# Complex Sphingolipid Profiling and Identification of an Inositol Phosphorylceramide Synthase in *Dictyostelium discoideum*

**DOI:** 10.1101/2023.07.07.548115

**Authors:** Stevanus A. Listian, Matthijs Kol, Edwin Ufelmann, Sebastian Eising, Florian Fröhlich, Stefan Walter, Joost C. M. Holthuis, Caroline Barisch

## Abstract

*Dictyostelium discoideum* is a professional phagocyte frequently used as experimental model to study cellular processes underlying the recognition, engulfment and infection course of microbial pathogens. Sphingolipids are abundant components of the plasma membrane that bind cholesterol, control vital membrane properties, participate in signal transmission and serve as adhesion molecules in recognition processes relevant to immunity and infection. While the pathway of sphingolipid biosynthesis has been well characterized in plants, animals and fungi, the identity of sphingolipids produced in *D. discoideum*, an organism at the crossroads between uni- and multicellular life, is not known. Combining lipidomics with a bioinformatics-based cloning strategy for key sphingolipid biosynthetic enzymes, we show here that *D. discoideum* produces phosphoinositol-containing sphingolipids with predominantly phytoceramide backbones. Cell-free expression of candidate inositol-phosphorylceramide (IPC) synthases from *D. discoideum* in defined lipid environments enabled identification of an enzyme that selectively catalyses the transfer of phosphoinositol from phosphatidylinositol onto ceramide. The corresponding IPC synthase, *Dd*IPCS1, is non-homologous to but shares multiple sequence motifs with yeast IPC and human sphingomyelin synthases and localizes to the Golgi apparatus as well as the contractile vacuole of *D. discoideum*. Collectively, these findings open up important opportunities for exploring a role of sphingolipids in phagocytosis and infection across major evolutionary boundaries.

## Introduction

Sphingolipids are a structurally diverse class of lipids with properties essential for the evolution of eukaryotic cells. They typically represent ∼ 15% of cellular lipids and are highly enriched in the plasma membrane (PM) where they modulate key physical membrane properties, including fluidity, thickness, and curvature (1, 2). Their affinity for cholesterol and ability to self-associate into laterally segregated membrane microdomains control a variety of membrane proteins including signaling receptors (3, 4). Besides their vital roles as structural components of cellular membranes, sphingolipids, and in particular intermediates of sphingolipid metabolism, function as signaling molecules that regulate a variety of biological processes, ranging from cell proliferation, cell death and cell migration to neurotransmission, angiogenesis and inflammation (5, 6). Sphingolipids also serve as receptor molecules for a plethora of extracellular ligands, thereby contributing to specific interactions between cells and their external environment (7). Additionally, various viruses and bacterial pathogens co-opt sphingolipids as receptors to gain entry into host cells (8) while several pore-forming toxins bind sphingolipids as receptors to perforate their target membrane.

The common feature of sphingolipids is a long-chain base (LCB), predominantly sphingosine in animals and phytosphingosine in plants and fungi (9, 10). N-acylation of the LCB with a long-chain fatty acid generates ceramide (Cer), a central intermediate of sphingolipid metabolism (11). Addition of a polar head group to the primary hydroxyl of Cer gives rise to complex sphingolipids. Based on their head group composition, complex sphingolipids can be grouped into phosphosphingolipids and glycosphingolipids. However, these categories are not mutually exclusive. Plants and fungi, for example, add phosphoinositol to phytoceramide to generate IPC, after which the inositol-containing head can be further decorated with one or more monosaccharides (12). While core features of the sphingolipid biosynthetic pathway and the subcellular organization of the underlying enzymatic machinery are conserved among eukaryotes (13, 14), organisms across different evolutionary branches generate complex sphingolipids with distinct polar head groups. For instance, plants and fungi generate IPC (10, 12) whereas mammals and nematodes produce phosphocholine-containing sphingomyelin (SM) (15–17). In contrast, insects like *Drosophila melanogaster* and some protozoa predominantly synthesize ethanolamine-phosphorylceramide (EPC) (18–23). Analogous to members of the lipid phosphate phosphatase (LPP) superfamily, known SM and IPC synthases are polytopic membrane proteins with an active site comprising a conserved catalytic triad of two histidine residues and one aspartate (24, 25). These residues participate in cleavage of the bond between the lipid hydroxyl and phosphate groups of the head group donor, phosphatidylinositol (PI) in the case of yeast IPC synthase, thus enabling the transfer of inositolphosphate from PI onto phytoceramide to produce IPC, releasing diacylglycerol as a side product. Animal SM synthases follow a similar LPP-type reaction cycle to produce SM, using phosphatidylcholine (PC) as head group donor. Besides sharing LPP-like sequence motifs, yeast IPC and animal SM synthases lack any obvious sequence homology (25).

The amoeba *D. discoideum* is an attractive model organism for cell biological research. Due to its relative simplicity and genetic tractability, it is particularly attractive for investigating fundamental processes including chemotaxis, phagocytosis, and autophagy. In infection biology, *D. discoideum* is frequently used as a surrogate host macrophage to investigate the pathogenesis of various microorganisms, including mycobacteria, *Legionella pneumophilla*, *Pseudomonas aeruginosa* and *Cryptococcus neoformans*. Apart from its importance for infection research, *D. discoideum* is also a versatile tool to study the cell biology of lipids. For example, recent work revealed that *D. discoideum* primarily utilises ether-linked inositol phospholipids for signalling (26) and that ether lipids play an important role during phagocytosis (27). We previously demonstrated that intracellular mycobacteria use *D. discoideum* triacylglycerols and phospholipids as major carbon and energy source (28, 29). Despite its wide acceptance as a model organism, the sphingolipidome of *D. discoideum* remains poorly characterized and many of the sphingolipid biosynthetic enzymes are yet to be identified. Considering that *D. discoideum* emerged after the plant-animal split and is evolutionary close to both *S. cerevisiae* and humans (30), elucidating the sphingolipid profile of this organism would offer valuable insights into the evolution of both sphingolipid biosynthesis and function.

Here, we conducted BLAST searches with sequences of previously identified sphingolipid biosynthetic enzymes to reconstruct the sphingolipid biosynthetic pathway in *D. discoideum*. As this approach did not yield any clue on the nature of complex sphingolipids in *D. discoideum*, we carried out mass spectrometry (MS)-based profiling of its sphingolipidome. This revealed that, analogous to plants and fungi, *D. discoideum* produces IPC. A bioinformatics-based search for candidate IPC synthases in *D. discoideum* coupled to their functional analysis in a cell-free expression system yielded a polytopic membrane protein, designated *Dd*IPCS1, which catalyses the transfer of phosphoinositol from PI onto ceramide to generate IPC. Localization studies showed that *Dd*IPCS1 resides both in the Golgi apparatus and contractile vacuole (CV) of *D. discoideum*.

## Material and Methods

### *D. discoideum* plasmids, strains and cell culture

All the *D. discoideum* material used in this study is listed in Supplemental Table 1. Wild type (AX2) was grown under static condition in a 10-cm dish at in 10 ml of HL5c medium (Formedium) supplemented with PS (100 U/ml penicillin and 100 mg/ml streptomycin).

To create CSS2-overexpressing cells, *css2* (DDB_G0268928) was amplified from *D. discoideum* cDNA and cloned into N- and C-terminal mCherry-fusion plasmids pDM1208 and pDM1210, respectively. The primers for cloning into pDM1208 were oMIB154 (5’-AGATCTATGGGAGTACAACAACAATCGG-3’) and oMIB155 (5’-ACTAGTCTATTTATTATTAAATT TTGATAAAATATTTTG-3’), while the primers for pDM1210 cloning were oMIB156 (5’-AGATCTAAAA TGGGAGTACAACAACAATCGG-3’) and oMIB157 (5’ ACTAGTTTTATTATTAAATTTTGATAAAA TATTTTG-3’). To generate the ZntD-GFP construct, ZntD was cleaved with SpeI and BglII from pDM1044-ZntD-mCherry (31) and cloned into pDM1045.

All plasmids used in this study are listed in Supplemental Table S1. Plasmids were electroporated into *D. discoideum* and selected with the appropriate antibiotic. Hygromycin was used at a concentration of 50 μg/ml, and neomycin (G418) at a concentration of 5 μg/ml.

### Lipidomics

To prepare cells for lipidomics, cells were passed in either defined SIH or complex HL5c medium (both from Formedium) for six days. Where indicated cells were treated additionally with 25 μM of AbA for 24 and 48 hrs, respectively. Confluent plates were harvested, washed and re-suspended in ice cold Sorensen buffer (SB). The cell suspensions were directly snap-frozen and stored at −80°C prior to extraction. For normalization, the protein concentration was determined using the Pierce BCA protein kit (Thermo Fisher Scientific).

For the LC/MS-MS lipidomics analysis, cell suspensions containing 200 μg of protein were mixed with 150 mM ammonium formate and subjected to lipid extraction using CHCl_3_/MeOH (2:1, v:v) following the protocol from Folch *et al.* (32, 33). Cer (18:0/17:1) was spiked to each sample as internal standard. Lipids were dried and subsequently dissolved with 50:50 mix of mobile phase A (60:40 water/acetonitrile, 10 mM ammonium formate, 0.1% formic acid) and mobile phase B (88:10:2 2-propanol/acetonitrile/H_2_O, 2 mM ammonium formate, 0.02% formic acid) before the analysis. For quantification, serial dilutions of LightSPLASH™ LIPIDOMIX® (Avanti Polar Lipids) were injected alongside the samples as external standards.

For the identification of IPCs and Cers, suspensions containing 60 μg of protein were subjected to lipid extraction and alkaline hydrolysis using dichloromethane/methanol (1:2, v:v) following a protocol described by Sullard *et al.* (34, 35). Also here, Cer (18:0/17:1) was added as internal standard. Subsequently, samples were mixed with 150 mM ammonium formate and organic solvents and mixed overnight at 48°C. Afterwards, 150 μl 1 M KOH were added and incubated for another 2 hrs at 37 °C under shaking conditions to hydrolyse glycerophospholipids. After neutralization with glacial acetic acid, the mixture was centrifuged and the supernatant was transferred to a new glass vial, dried, and re-suspended with a 65:35 mixture of mobile phase A and B (see above).

The HPLC run was performed using a C30 reverse-phase column (Thermo Acclaim C30, 2.1 250 mm, 3 mm, operated at 40°C; Thermo Fisher Scientific) connected to a Shimadzu LC-20AD HPLC system. A binary solvent system was used (mobile phases A and B) in a 20 min gradient at a flow rate of 300 μl/min. The HPLC was connected to a QExactivePLUS orbitrap mass spectrometer (Thermo Fisher Scientific) equipped with a heated electrospray ionization (HESI) probe. The maximum injection time for full scans was 100 ms with a target value of 3,000,000 at a resolution of 70,000 at m/z 200 and a mass range of 200 – 1200 m/z in positive and negative ion mode. The 5 and 10 most intense ions from the survey scan were selected and fragmented with high-energy collision dissociation with normalized collision energy of 25, 30, 35 (samples extracted with Folch) and 25, 30 (samples extracted with Sullards), respectively. Target values for MS/MS were set at 100,000 with a maximum injection time of 50 ms at a resolution of 35,000 at *m/z* 200.

For the quantification of total lipids, PIs, and ether/ester lipids, peaks were analysed using the Lipid Search algorithm (Thermo Fisher Scientific). Product ion (MS2) and precursor ion (MS1) peaks were defined from scanning through raw files. From the intensities of lipid standards, the absolute values for each lipid classes in mol % were calculated. SM, PC, LPC, DAG, TAG and Cer were acquired in the positive ion mode, while PE, PI, PG, PS and LPE were acquired in negative ion mode. Candidate molecular species were identified by a database search for positive (+H+; +NH4+) or negative ion adducts (-H-; +COOH-). Mass tolerance was set to 20 ppm and 5 ppm for the precursor mass from the positive ion mode and the negative ion mode, respectively. For the peak alignment between samples, the retention time (RT) tolerance were set to 0.5 and 0.21 min for the positive and negative ion mode, respectively. Samples were aligned within a time window and results were combined in a single report.

For the quantification of IPC from samples extracted by Folch extraction, the software FreeStyle™ 1.8 SP2 (Thermo Fisher Scientific) was used. To this end, the peak area of precursor ions normalized to the internal standard were calculated. For the quantification of IPC and Cer from samples extracted by Sullards, the peaks were analysed with Skyline (36, 37). The top thirteen most abundant IPC species were selected. To confirm that the quantified peak represents IPC, the presence of MS/MS fragments characteristic to IPC (m/z 241 representing the inositol-1,2-cyclic phosphate anion and m/z 259 representing the inositol-monophosphate anion) in all IPC species was checked (38). The peak area of precursor ions from each IPC species was normalized against the total IPC signal from each sample. The complete dataset of the lipidomics analysis is shown in Supplementary Table 2.

#### Preparation of liposomes

Liposomes were prepared as previously described (39). Briefly, phospholipid stocks were prepared in CHCl_3_:MeOH (9:1, v:v) and stored at −20 °C. The phospholipid concentration was determined following (40). Using a mini extruder (Avanti Polar Lipids), unilamellar liposomes were produced from a defined lipid mixture as stated in the figure legends: egg PC: DOPE: wheat germ PI 2:2:1 mol %, egg PC: DOPE 1:1 mol %; egg PC: wheat germ PI 1:1 mol %; and DOPE: wheat germ PI 1:1 mol %. To create a thin lipid film, 20 μmol of total lipids were dried with nitrogen gas. Dried lipids were re-suspended in 1 ml lipid rehydration buffer (25 mM HEPES (pH 7.5) and 100 mM NaCl) to create a 20 mM lipid suspension. After six freeze-thaw cycles, liposomes were extruded through 0.4 μM and 0.1 μM polycarbonate membrane filter (Whatman-Nucleopore) to obtain an average diameter of 100 nm. Liposome suspensions were snap-frozen and stored at −80 °C.

#### Cell-free expression (CFE) and CSS activity assay

The CFE expression of CSS was performed as previously described (39). Briefly, proteinase K was used to remove trace amounts from RNAse from pEU-Flexi-CSS1, pEU-Flexi-CSS2 and pEU-Flexi-SMS2 expression constructs (Supplemental Table 1). CSS1 and CSS2 were synthesised together with a V5-tag using GeneArt DNA Synthesis (Thermo Fisher Scientific) and then cloned into pEU-Flexi. After phenol/CHCl_3_ extraction, the constructs were dissolved in water at 1.8 μg/μl. The mRNA was transcribed in a 120 μl reaction volume containing 21.6 μg of DNA; 2 mM each of ATP, GTP, CTP, and UTP; 45.6 units of SP6 RNA polymerase; 96 units of RNase inhibitor in 80 mM HEPES-KOH (pH 7.8); 20 mM of Mg-acetate; 2 mM of spermidine hydrochloride and 10 mM of DTT (41). After a 4 hrs incubation at 37°C and a centrifugation at 3,400 g for 5 min at RT, the supernatant was harvested and subsequently used for the cell-free translation reaction. Each 100 μl translation reaction contained: 2 mM of liposomes, 40 μg/ml of creatine kinase, 15 OD_260_ of WEPRO2240 WGE, 0.3 mM each of all 20 amino acids, 5 mM of HEPES-KOH (pH 7.8), 50 mM of potassium acetate, 1.25 mM of Mg-acetate, 0.2 mM of spermidine, 0.6 mM of ATP, 125 μM of GTP, 8 mM of creatine phosphate and 20 μl of the mRNA-containing supernatant. The translation mixture was then incubated for 4 hrs at 26 °C. The translation products were directly used for the CSS assay. Protein expression was confirmed using western blots and anti-V5.

For the CSS activity assay, the reaction volume was set to 100 μl: 50 μl of the CFE product were incubated first with 5 μM of NBD-Cer, 25 mM of HEPES (pH 7.4), 75 mM of NaCl and 1 mM of MgCl_2_ for 10 min on ice. This was followed by a 60 min incubation at 22°C under shaking condition. When indicated, Aba was added, ethanol served as vehicle control. The reaction was terminated by adding 375 μL of CHCl_3_:MeOH (1:2, v:v) and stored at −20°C prior to lipid extraction and TLC analysis. Where indicated yeast lysates served as control. The preparation of the post-nuclear supernatant fraction of *S. cerevisiae* was performed as described previously (39).

### Live fluorescence microscopy

To monitor the subcellular distribution of CSS2-mCherry or mCherry-CSS2 by live imaging, cells were transferred to either 4- or 8-well µ-slides (ibidi) and imaged in low fluorescent medium (LoFlo, Foremedium) with a Zeiss Cell observer spinning disc (SD) microscope using the 63x oil objective (NA 1.46). Images were further processed and analysed with ImageJ.

### Immunofluorescence

The anti-vatA, anti-vacA, anti-p80 and anti-Golgi antibodies were obtained from the Geneva antibody facility (Geneva, Switzerland). The anti-PDI antibody was provided by Markus Maniak (University of Kassel, Germany). As secondary antibodies, anti-mouse and anti-human IgG coupled to CF488R (Biotium) or CF640R (Biotium) were used.

To prepare cells for IF, *D. discoideum* was either fixed with 4 % PFA/picric acid or with cold MeOH as described previously (42). Where indicated, cells were incubated overnight with 3 μm latex beads (Sigma Aldrich) before the fixation. Cells were embedded using ProlongGold antifade with DAPI (Molecular Probes). Images were recorded with an Olympus LSM FV-3000 confocal microscope and the 60x PLAPON-SC 1.4 NA oil immersion objective. To improve signal-to-noise, indicated images were deconvolved using Huygens Software from Scientific Volume Imaging (Netherlands). The images were further processed and analysed with ImageJ and Inkscape.

### SDS-PAGE and western blotting

For western blotting of mCherry-tagged CSS2, cells were washed, harvested and incubated with 2x Laemmli buffer. For western blotting of V5-tagged proteins, samples were added to urea buffer containing 5% ß-mercaptoethanol instead. After SDS-PAGE, proteins were transferred for 50 min at 120 V to a nitrocellulose membrane (Amersham^TM^ Protran^TM^, Premium 0,45 µm NC). To check the efficiency, Ponceau S staining was performed. For the detection of mCherry-tagged proteins, the membranes were blocked and incubated with an anti-mCherry primary (Rockland) and a goat anti-rabbit secondary antibody coupled to horseradish peroxidase (HRP) (BioRad). For V5-tagged proteins, an anti-V5 primary (Invitrogen; 1:4000) and a goat anti-mouse secondary antibody coupled to horseradish peroxidase (HRP) (BioRad, 1:4000) were used. The detection of HRP was accomplished using the Pierce^TM^ ECL Western Blotting Substrate (Thermo Scientific).

### Selection and topology prediction of D. discoideum CSS

The *D. discoideum* AX4 proteome was downloaded from UniProt (ID: UP000002195) and searched using Prosite (https://prosite.expasy.org/) for the presence of the H-X3-D-X3-[GA]-X3-[GSTA] motif. The FASTA files of all 52 protein hits were compiled and submitted as queries to TMHMM 2.0 to search for the presence of TMDs (43). Proteins with >3 TMDs were selected for further screening. Among the seven remaining candidates, proteins with known biochemical function (with annotation at dictybase.org or identification by BLAST search) were excluded yielding two potential CSS candidates.

AlphaFold was used to predict the membrane topology of *Dd*CSS1 and *Dd*CSS2 (44). MAFFT version 7 algorithm (https://mafft.cbrc.jp/) was used to construct phylogenetic tree and protein sequences (45). The following parameters was adopted for MAFFT alignment: G-INS-i strategy, unalignlevel 0.0 and “try to align gappy regions away”. NJ conserved sites and the JTT substitution model was used to generate a phylogenetic tree in phylo.ilo. Numbers on the branches indicate bootstrap support for nodes from 100 bootstrap replicates.

## Results

### Reconstruction of the sphingolipid biosynthetic pathway in D. discoideum

While sphingolipids are ubiquitous among eukaryotes, the pathway of *de novo* sphingolipid biosynthesis in *D. discoideum* has remained largely unexplored. This led us to search the *D. discoideum* database for homologs of key sphingolipid biosynthetic enzymes previously identified in *A. thaliana* (46), *S. cerevisiae* (47) and *H. sapiens* (48)(Fig.1, Supplemental Table 3). The first committed and rate-limiting step of sphingolipid synthesis is the condensation of serine and palmitoyl-CoA, a reaction catalysed by serine-palmitoyltransferase (SPT) yielding ketodihydrosphingosine or 3-ketosphinanine (3-KDS) (Fig. 1). In eukaryotes, SPT comprises a core heterodimer of two large catalytic subunits and two accessory small subunits (ssSPT). While the genome of *D. discoideum* encodes homologues of the two large catalytic subunits (SptA and SptB; Table 1), we were unable to identify any ssSPT homologue. SPT forms a so-called SPOTS complex with the regulatory Orosomucoid (Orm) proteins and the phosphatidylinositol-4-phosphate phosphatase Sac1. *D. discoideum* appears to have only one Orm-like protein (DDB_G0288847), we found two structural homologues of Sac1 (Sac1 and Sac1-like). A recent study revealed the cryo-EM structure of the Cer-bound SPOTS complex in yeast and provided evidence for the presence of a Sac1-containing SPOTS complex in *D. discoideum* (47). SPT gives rise to 3-KDS, which is then further reduced to sphinganine (dihydropshingosine) by 3-keto-sphinganine reductase (Ksr). In *D. discoideum*, there are two identical copies of this enzyme in AX3 and AX4 (but not AX2), known as KsrA-1 and KsrA-2. Sphinganines are then converted into dihydroceramides by Cer synthases (CerS) through the addition of fatty acid moieties. Mammals, plants and yeast each contain three to six CerS isoforms that produce dihydroceramides with distinct acyl chain lengths (49–51). This is in sharp contrast to *D. discoideum,* which contains only one CerS homologue (CrsA)(52). Dihydrosphingosines and dihydroceramides are generally further modified by desaturases and hydroxylases. A BLAST search for *D. discoideum* homologues of the yeast sphinganine C4-hydroxylase Sur2 produced three hits, here referred to as lipid hydroxylase LhsA, LhsB and LhsC. Sphingolipid desaturases are present in human (53) and *A. thaliana* (54, 55) but absent in *S. cerevisiae* (56). In *D. discoideum* we found one dihydroceramide desaturase homologue (DesA). The amide-linked acyl chain in sphingolipids usually undergoes further hydroxylations. A search for *D. discoideum* homologues of the yeast Cer fatty acid hydroxylase Scs7 and the *A. thaliana* Cer fatty acid hydroxylases FAH1 and FAH2 in each case yielded one hit, namely LhsD.

**Fig. 1.**
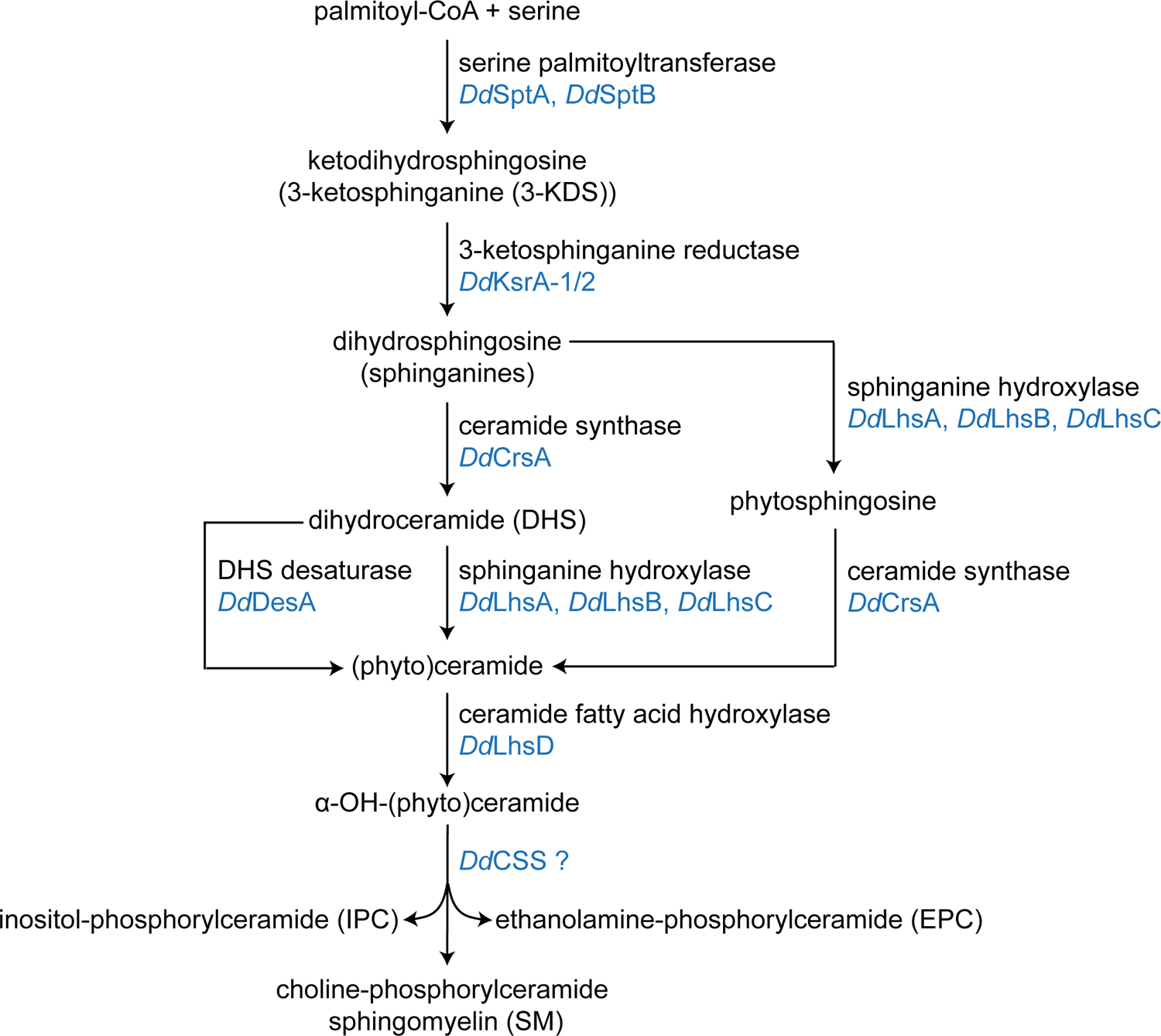
The sphingolipid biosynthetic pathway in *D. discoideum*. The pathway was reconstructed based on BLAST searches for *D. discoideum* homologues (marked in blue) of key sphingolipid biosynthetic enzymes identified in *S. cerevisiae*, *A. thaliana* and *H. sapiens* (Supplemental Table 3). Note that this approach did not yield any *D. discoideum* homologues of complex sphingolipid synthases (CSS) responsible for the production of inositol-phosphorylceramide (IPC), ethanolamine-phosphorylceramode (EPC) or sphingo-myelin (SM) in fungi, plants or animals.

**Table 1.** Sphingolipid biosynthetic enzymes in *D. discoideum*, *A. thaliana*, *S. cerevisiae* and *H. sapiens*.

| Enzyme | *D. discoideum* | *A. thaliana* | *S. cerevisiae* | *H. sapiens* |
| --- | --- | --- | --- | --- |
| serine palmitoyltransferase<br>(large subunits) | SptA (DDB_G0268056),<br>SptB (DDB_G0291283) | LCB1, LCB2a,<br>LCB2b | Lcb1, Lcb2 | SPTLC1,<br>SPTLC2,<br>SPTLC3 |
| 3-ketodihydrosphingosine<br>reductase | KsrA-1 (DDB_G0274015),<br>KsrA-2 (DDB_G0272883) | KSR1, KSR2 | Tsc10 | KDSR |
| sphingolipid desaturase | DesA (DDB_G0269738) | SLD1, SLD2,<br>DES1 | non-existent | DEGS1 |
| ceramide synthase | CrsA (DDB_G0282607) | LOH1, LOH2,<br>LOH3 | Lac1, Lag1,<br>Lip1 | CERS1, CERS2,<br>CERS3, CERS4,<br>CERS5, CERS6 |
| sphingolipid hydroxylase | LhsA (DDB_G0269788),<br>LhsB (DDB_G0270946),<br>LhsC (DDB_G0279611) | SBH1, SBH2 | Sur2 | DES2 |
| ceramide acyl hydroxylase | LhsD (DDB_G0269908) | FAH1, FAH2 | Scs7 | non-existent |

Collectively, these results indicate that *D. discoideum* has the capacity to produce Cers with a desaturated LCB and carrying hydroxyl groups in both the LCB and amide-linked acyl chain. Cers can be further derivatized by addition of a headgroup at C1 to generate complex sphingolipids like glucosylceramide (GlcCer), the precursor of complex glycosphingolipids that contain a few to dozens of sugar residues. Alternatively, the C1 of Cer can be modified with a phospho-alcohol to generate SM, EPC or IPC. A search for *D. discoideum* homologues of GlcCer synthases in *Homo sapiens* (*Hs*CEGT), *Candida albicans* (*Ca*CEGT) or *A. thaliana* (*At*CEGT) did not produce any hits. Likewise, BLAST searches using sequences of the enzymes responsible for the production of SM (*Hs*SMS1), EPC (*Hs*SMSr) or IPC (*Sc*Aur1, *At*IPCS1) did not yield any homologues in *D. discoideum.* Consequently, our homology searches did not yield further insights into the nature of the complex sphingolipids produced by *D. discoideum*.

### Phosphatidylethanolamine is the most abundant phospholipid in D. discoideum

The synthesis of complex phosphosphingolipids involves the transfer of the polar headgroup from a phospholipid donor such as phosphatidylcholine (PC), phosphatidylethanolamine (PE) or PI onto C1 of Cer. Interestingly, eukaryotic organisms evolved different head group dominances regarding their phospholipid and sphingolipid production. Mammals contain more PC than PE or PI and predominantly synthesize phosphocholine-containing SM (57). In contrast, plants and yeast contain PI as their most abundant glycerophospholipid and produce phosphoinositol-containing sphingolipids (58, 59) whereas insects synthesize phosphoethanol-based glycerophospholipids and sphingolipids (18, 22). Thus, the insect lipidome appears to be ethanolamine-centric, in contrast to choline-centric mammals and inositol-centric plants and yeast. To determine the head group dominance in *D. discoideum*, we performed an untargeted LC-MS/MS analysis of total lipid extracts from *D. discoideum* cultured in defined medium (SIH, Formedium). We found that PE is by far the most prevalent lipid class in *D. discoideum,* representing ∼ 70 mol % of all lipids analysed, with PC and PI representing ∼ 10 mol % and ∼ 7 mol %, respectively (Fig. 2A, Supplemental Table 2). These data are consistent with those reported previously by others (27, 60, 61). Besides, diacylphospholipids (∼ 88.7 mol %), *D. discoideum* contains substantial amounts of lysophospholipids (∼ 6.8 mol %), notably lysophosphatidyl-ethanolamine (LPE), Cers (∼ 0.6 mol %), diacylglycerol (DAG, ∼ 2.3 mol %) and triacylglycerols (TAG; ∼ 1.6 mol %) (Fig. 2A).

**Fig. 2.**
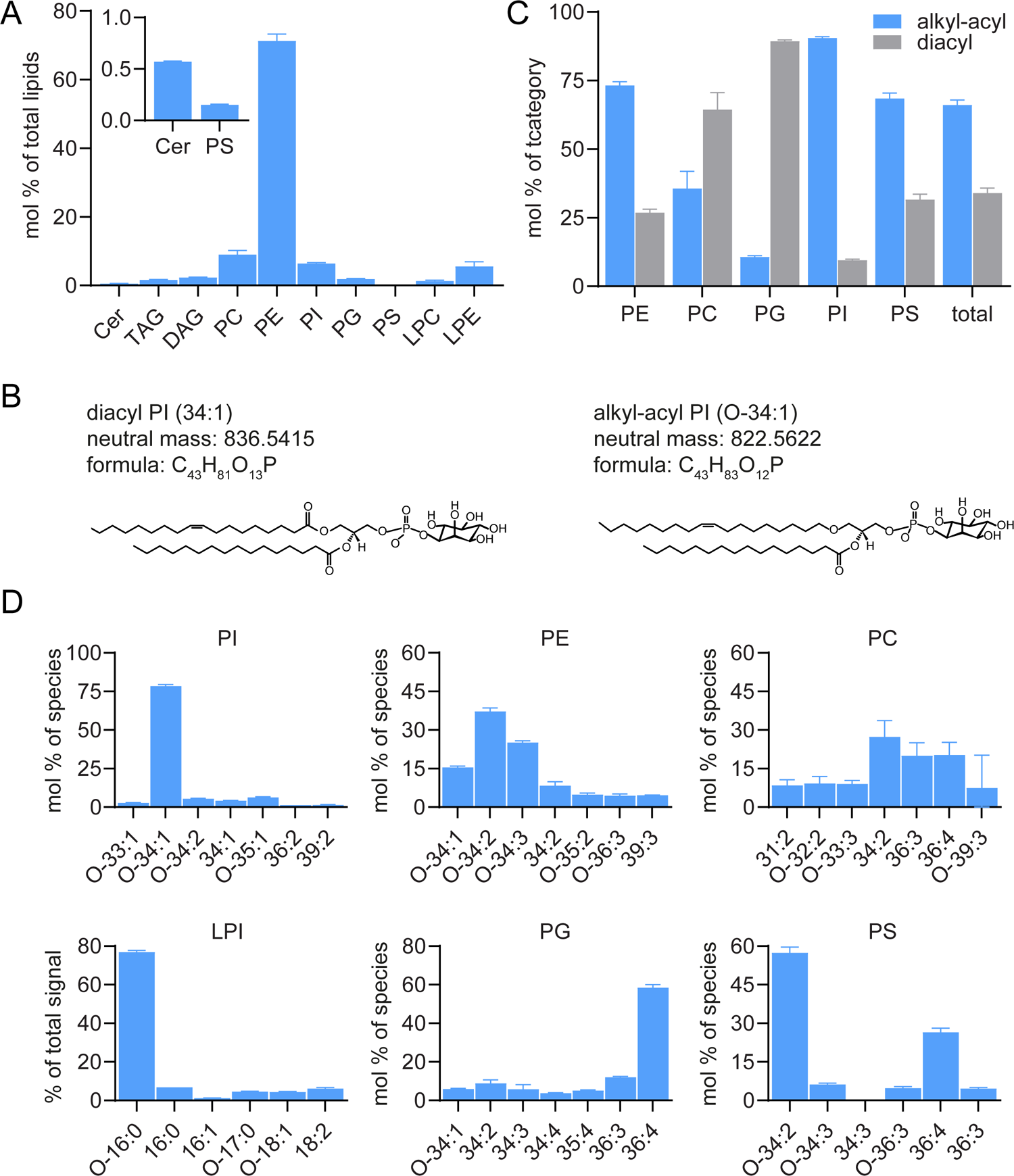
Quantitative analysis of the main lipid classes in *D. discoideum*. A: Levels of indicated lipid classes in total lipid extracts from *D. discoideum* were determined by LC-MS/MS and expressed as mol % of total lipid analysed. Note that PE is the dominant phospholipid in *D. discoideum*. Inset shows a close-up of ceramide (Cer) and PS levels. B: Molecular structures of ester (diacyl) PI (34:1) and ether (alkyl-acyl) PI (O-34:1). C: Quantification of the distribution of alkyl-acyl species (blue) and diacyl species (grey) within each phospholipid class. D: Lipid molecular species composition within each phospholipid class. Shown are the seven most abundant species of each category, except for PS and LPI where only six lipid species were detected. Lipids were quantified as mol % of total lipids analysed, except for LPI. LPI was quantified as % of total signal due to the absence of an LPI standard. *D. discoideum* was grown for six days in SIH (defined) medium prior to lipid extraction. Lipids were extracted following (32). Data are means ± SD (n=3).

Phospholipids have either an ester or an ether bond at the *sn*-1 position. Figure 2B provides molecular structures of an ester lipid (diacyl PI) and an ether lipid (alkyl-acyl PI). In *D. discoideum*, the majority of inositol phospholipids are ether lipids with a novel plasmanyl-O-34:1 structure (with the “O” indicating ether linkage of one carbon chain to the glycerol) (26). Strikingly, this also accounts for all other *D. discoideum* phospholipids, except for PC and PG, in which diacyl bonds are more abundant (Fig. 2C). In addition, we found that the composition of PI species was highly similar to that reported by a previous study (26) (Fig. 2D). In the same study, Clark *et al*. reported that PI (O-34:1) is comprised of an ether-linked C16:0 fatty acid at its *sn*-1 position. This is consistent with the fact that in our hands lysophosphatidyl-inositol (LPI) (O-16:0) is the most abundant LPI species (Fig. 2D).

We conclude that PE is the most abundant phospholipid in *D. discoideum* and that the majority of *D. discoideum* phospholipids are ether lipids that contain alkyl-acyl bonds.

### D. discoideum produces phosphoinositol-containing sphingolipids

The high abundance of PE implies that *D. discoideum* might use PE as headgroup donor to synthesis EPC as previously demonstrated in *Drosophila* and other insects (18, 62). To address this possibility and identify complex sphingolipids in *D. discoideum* by lipidomics, we used a lipid extraction protocol optimized for the isolation of complex sphingolipids (34). In brief, total lipids were extracted with dichloromethane/methanol and subjected to alkaline hydrolysis to deacylate the phospholipids and hence eliminate isobaric overlaps between sphingolipids and phospholipids. The efficiency of this protocol is reflected by the absence of PI (O-34:1) after alkaline hydrolysis (Supplemental Fig. S1). To identify complex sphingolipids, we performed a targeted search using the LipidCreator workbench from Skyline (37). This enabled the detection of various species of IPC, with IPC (38:0;4) being the most abundant IPC species in *D. discoideum* (Fig. 3A-B). The identity of IPC (38:0;4) was supported by the presence of MS2 fragments (product ions) characteristic for IPC (inositol-1,2-cyclic phosphate anion and inositol monophosphate anion at *m/z* 241 and 259, respectively) (Fig. 3A, inset). In addition, the product ion at [M-H-180]^-^ represents the loss of the inositol residue and corroborates the claim that the lipid described in Figure 3A is indeed IPC. However, using our current approach, we were not able to conclusively resolve the length of the N-linked acyl chain and LCB from the IPC (38:0;4) peaks. We therefore analysed the product ion fragmentation of Cer (38:0;4), which is presumably the precursor of IPC (38:0;4) (Supplemental Fig. S2A). The product ions representing LCB (18:0;3) and LCB (20:0;3) were both detected in the precursor peak of Cer (38:0;4) (Supplemental Fig. S2B). However, the intensity of LCB (18:0;3) was much higher than the one of LCB (20:0;3). This suggests that Cer (18:0;3/20:0;1) constitutes the major proportion of Cer (38:0;4), while Cer (20:0;3/18:0;1) accounts for the minority fraction. From this we infer that the most abundant IPC species in *D. discoideum* is IPC (18:0;3/20:0;1).

**Fig. 3.**
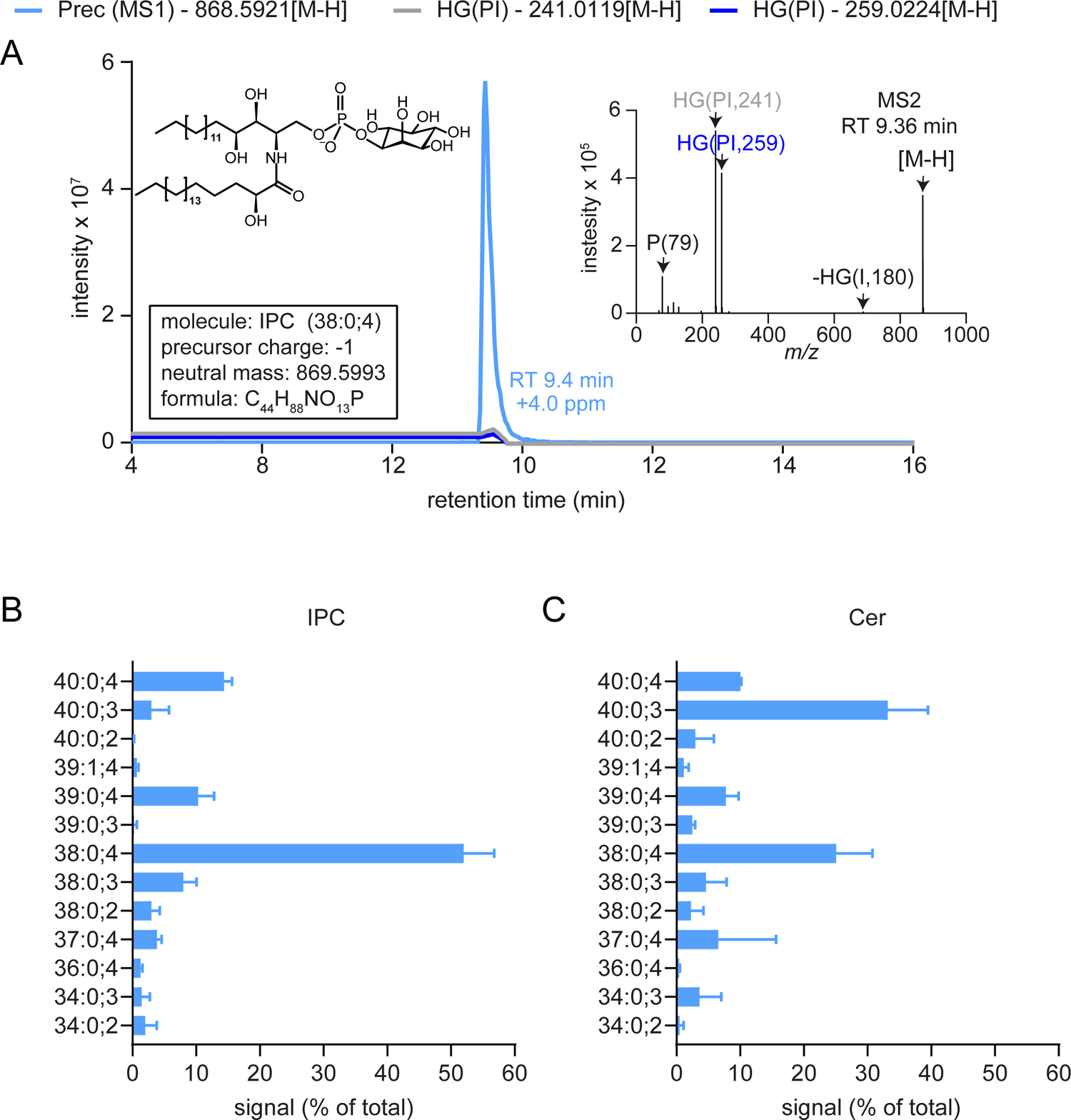
Profiling of IPC and ceramide (Cer) species in *D. discoideum*. A: Detection of IPC (38:0;4) by LC-MS/MS. The inset shows the product ion fragmentation of IPC (38:0;4) at RT 9.36 min. B: LC-MS/MS analysis of the main IPC species in *D. discoideum.* C: LC-MS/MS analysis of the Cer species in *D. discoideum* that correspond to the IPC species shown in (B) with respect to chain length, hydroxylation status and saturation level. *D. discoideum* was grown in HL5c (complex) medium and then subjected to total lipid extraction according to (34). The peak area of the precursor ions ([M-H]) from each IPC and Cer species was quantified and normalized with the total peak area from all IPC/Cer species using Skyline. Data are means ±SD (n=3).

Besides IPC (38:0;4) (∼ 52 % of total IPC), *D. discoideum* contains various other IPC species, of which the most abundant are IPC (40:0;4, ∼ 14.3 %), IPC (39:0;4, ∼ 10.3 %) and IPC (38:0;3, ∼ 8.0 %; Fig. 3B). The most abundant ceramide species are Cer (40:0;3) (∼ 35.9 % of total ceramide species), followed by Cer (38:0;4) (∼ 27.3 %), Cer (40:0;4) (∼11.0 %) and Cer (39:0;4) (∼ 8.5 %; Fig. 3C). This indicates that, like in fungi and plants, complex sphingolipids in *D. discoideum* are mainly based on “phytoceramides” that contain three or more hydroxyl groups. Our analysis also revealed the presence of minor amounts of unsaturated IPC and Cer species as well as odd chain IPCs and Cers (Fig. 3B and 3C).

Given that *D. discoideum* contains more PE and PC than PI (Fig. 2A), we also analysed the alkaline-hydrolysed *D. discoideum* lipid extracts for the presence of EPC or SM species. However, we did not detect any ethanolamine or choline-containing counterparts of the most abundant IPC species (Fig. 4; Supplemental Fig. S3). From this we conclude that, analogous to plants and yeast, *D. discoideum* synthesises IPC. We next set out to identify the corresponding IPC synthase.

**Fig. 4.**
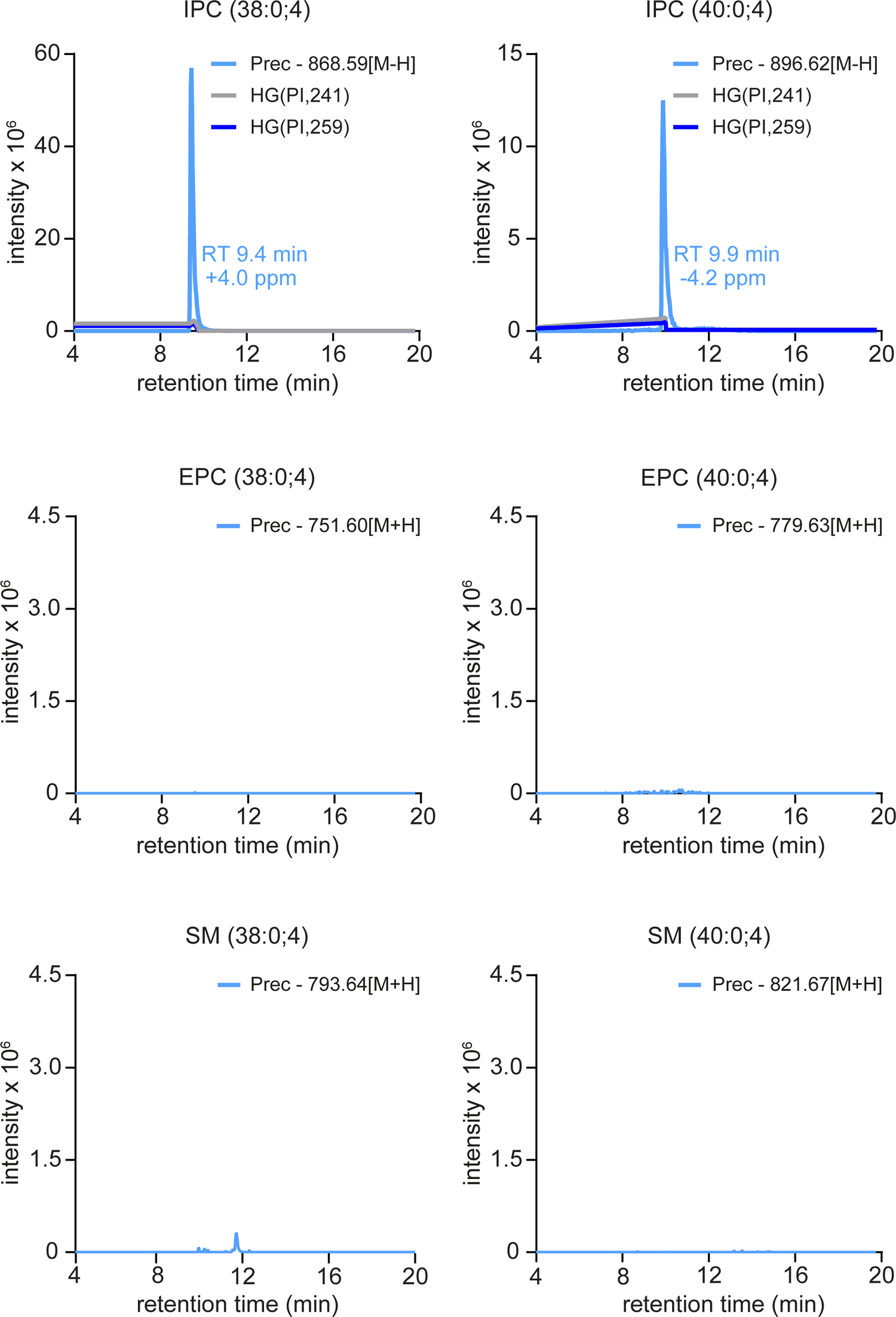
*D. discoideum* exclusively produces inositol-containing phosphosphingolipids. Detection of IPC (38:0;4) and IPC (40:0;4) but not their SM or EPC counterparts by LC-MS/MS analysis in alkaline-treated lipid extracts from *D. discoideum*. *D. discoideum* was grown in HL5c (complex) medium. Lipids were extracted following (34). Data shown are representative of three independent experiments (n=3).

### A bioinformatics-based search for IPC synthase in D. discoideum

As *D. discoideum* lacks structural homologues of the IPC synthases present in plants and yeast, we pursued a bioinformatics and functional cloning strategy to identify the IPC synthase(s) in *D. discoideum.* A similar strategy previously led to identification of the human SM synthases SMS1 and SMS2 (25). As outlined in Fig. 5A, the selection criteria for candidate IPC synthases in *D. discoideum* are: (1) presence of a short sequence motif (H-X3-D-X3-[GA]-X3-[GSTA]) shared among all known lipid phosphate phosphatases (LPPs) and complex sphingolipid synthases (CSSs), including human SMS1/2 and the yeast IPC synthase *Sc*Aur1p; (2) presence of multiple (>3) transmembrane domains (TMDs) given that all known CSSs have six to eight predicted TMDs; (3) biochemical function should be unknown; and (4) the domain architecture should be similar to known CSSs. Of the 12,734 protein sequences analysed, two met all four selection criteria and therefore qualified as candidate *D. discoideum* IPC synthases, which we designated *Dd*CSS1 and *Dd*CSS2 (Fig. 5A). A phylogenetic analysis of CSS sequences from humans, fungi, parasites, plants and *D. discoideum* revealed that *Dd*CSS1 is more closely related to human SMS and IPCS from plants and protozoa, while *Dd*CSS2 is more closely related to fungal IPC synthases (Fig. 5B).

**Fig. 5.**
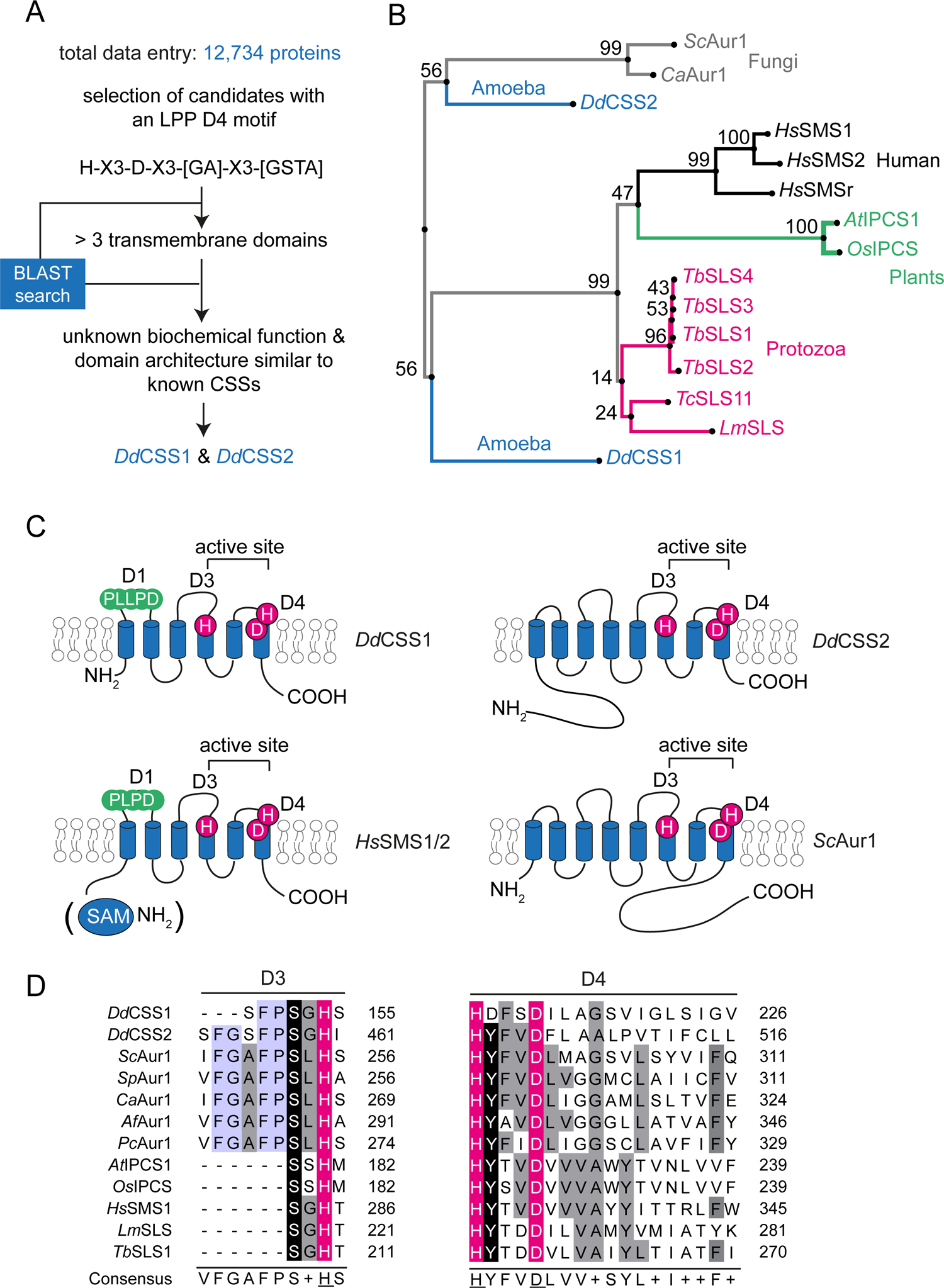
Selection and phylogenetic analysis of candidate IPC synthases in *D. discoideum.* A: Search of the *D. discoideum* proteome for polytopic membrane proteins that contain the LPP-like sequence motif H-X3-D-X3-[GA]-X3-[GSTA] and meet additional selection criteria as indicated yielded two candidate complex sphingolipid synthases (CSS): *Dd*CSS1 (DDB_G0284367) and *Dd*CSS2 (DDB_G0268928). B: Phylogenetic tree of *Dd*CSS1, *Dd*CSS2 and IPC/SM synthases from different species: *Sc*Aur1 from *Saccharomyces cerevisiae* (UniProt accession: P36107); *Ca*Aur1 from *Candida albicans* (O13332); *At*IPCS1 (Q9M325) from *Arabidopsis thaliana*; *Os*IPCS1 from *Oryza sativa* (Q5N7A7); *Hs*SMS1, *Hs*SMS2 and *Hs*SMSr (Q86VZ5, Q8NHU3 and Q96LT4) from *Homo sapiens*; *Lm*SLS (E9AFX2) from *Leishmania*; *Tb*SLS1-4 (Q38E54, Q38E56, Q38E53 and Q38E55) from *Trypanosoma brucei; Tc*SLS11 from *Trypanosoma cruzi* (UniProt accession: Q4E4I4). Protein sequences were aligned with MAFFT (https://mafft.cbrc.jp) using the G-INS-i strategy, unalignlevel 0.0 and “try to align gappy regions away” to generate a phylogenetic tree in phylo.ilo using NJ conserved sites and the JTT substitution model. Numbers on the branches indicate bootstrap support for nodes from 100 bootstrap replicates. C: Membrane topology of *Dd*CSS1 and *Dd*CSS2 reconstructed by AlphaFold (https://alphafold.ebi.ac.uk/)(Supplemental Fig. 4C). D: Alignment of D3 and D4 sequence motifs in *Dd*CSS1, *Dd*CSS2 and known IPC/SM synthases from different organisms. *Sp*Aur1 (Q10142) from *Schizosaccharomyces pombe*; *Af*Aur1 (Q9Y745) from *Aspergillus fumigatus*; *Pc*Aur1 (Q9Y745) from *Pneumocystis carinii* (Q6AHV1). Alignment was performed with Jalview workbench (87) using the in-built MAFFT alignment option (E-INS-i). Identical residues are shaded in black, conservative residue substitutions are shaded in grey and conserved residues that are part of the catalytic triad are shaded in magenta. Residues conserved among fungal IPC synthases, *Dd*CSS1 and *Dd*CSS2 are shaded in blue.

Sequence analysis of *Dd*CSS1 and *Dd*CSS2 revealed two conserved sequence motifs which correspond to the D3 and D4 motifs in human SMS1/2 (25). The D3 (C-G-D-X3-S-G-<u>H</u>-T) and D4 (<u>H</u>-Y-T-X-<u>D</u>-V-X3-Y-X6-F-X2-Y-H) motifs are similar to the C2 and C3 motifs in LPPs (63) and include the histidine and aspartate residues (magenta) that form the catalytic triad mediating the nucleophilic attack on the lipid phosphate ester bond (Fig. 5C-D)(64). Unlike *Dd*CSS2, *Dd*CSS1 contains a D1 (P-L-P-D)-like motif (“P-L-L-P-D”) in its first exoplasmic loop (Fig. 5C, Supplemental Fig. S4A) that is also present in human SMS1/2 (25) and protozoan IPC synthases (65). Neither *Dd*CSS1 nor *Dd*CSS2 contains the D2 (R-R-X8-Y-X2-R-X6-T) motif that is typically located in the third transmembrane span of SMS family members. The Aur1-related IPC synthases in fungi contain two unique sequence motifs, here designated IPCS1 (D-h-h-n-W-X2-Y-X3-H-X3-P) and IPCS2 (Y-X3-G-X3-G-L-X-R-X-D)(24). The tyrosine residue at position 254 (Y) in the IPCS1 motif and the peptide sequence R-I-D at position 444-446 in the IPCS2 motif are conserved in *Dd*CSS2 but not in *Dd*CSS1 (Supplemental Fig. S4B).

Using models derived from AlphaFold (Supplemental Fig. S4C), *Dd*CSS1 is predicted to contain six membrane-spanning alpha helices, hence analogous to human SMS1/2 and plant IPCS (Fig. 5C). In contrast, *Dd*CSS2 is predicted to contain eight transmembrane domains, analogous to *Sc*Aur1. According to the structures, both N- and C-termini of *Dd*CSS1 and *Dd*CSS2 would be cytosolic while their active site would be positioned on the exoplasmic leaflet (Fig. 5C). This is in line with the experimentally established membrane topologies of SMS and LPP family members (25).

In conclusion, the membrane topology and location of the catalytic triad in *Dd*CSS1 and *Dd*CSS2 indicate that both proteins qualify as candidate CSS. Moreover, *Dd*CSS2 shares additional structural features with IPC synthases in fungi.

### DdCSS2 displays IPC synthase activity

To determine whether *Dd*CSS1 and *Dd*CSS2 possess IPC synthase activity, the corresponding mRNAs were synthesized *in vitro* and translated in a wheat germ extract (WGE) in the presence of unilamellar liposomes prepared from a mixture of PC, PE and PI (Fig. 6A). This approach previously enabled the cell-free production of enzymatically active *Hs*SMS1/2 (39) and protozoan IPC synthases (23). In brief, the cDNAs of *Dd*CSS1 and *Dd*CSS2 were cloned downstream of the Sp6 promoter in pEU Flexi-vector pFLx (41). A pFLx construct encoding *Hs*SMS2 served as control. To facilitate detection of the cell-free-produced proteins, a V5 epitope was introduced at their C-termini. As shown in Fig. 6B (top panel), cell-free translation of *Dd*CSS1/2 and *Hs*SMS2 mRNA in the presence of liposomes in each case yielded a V5-tagged protein of the expected size. No such protein was detected when translation was performed in the absence of mRNA. Next, proteoliposomes were incubated with the fluorescent short-chain (C6) Cer analog NBD-Cer and the formation of complex NBD-labelled sphingolipids was monitored by TLC. As expected, proteoliposomes containing *Hs*SMS2 supported production of NBD-SM (Fig. 6B, bottom panel; Supplemental Fig. S5A). In contrast, proteoliposomes containing *Dd*CSS1 did not yield any obvious NBD-labelled reaction product. However, the reaction with *Dd*CSS2 proteoliposomes yielded an NBD-labelled product with a retention value distinct from NBD-SM but similar to that of NBD-IPC (Fig. 6B, bottom panel; Supplemental Fig. S5A). Importantly, omission of PI from *Dd*CSS2 proteoliposomes abolished production of the NBD-labelled lipid while omission of PC or PE did not (Fig. 6C, bottom panel; Supplemental Fig. S5B-C). Conversely, omission of PC from *Hs*SMS2 proteoliposomes diminished but did not entirely eliminate production of NBD-SM, presumably because the WGE contains residual amounts of PC (Supplemental Fig. S5D). From this we conclude that *Dd*CSS2 possesses IPC synthase activity. The activity of *Dd*CSS1 remains to be established. To address whether *Dd*CSS2 is responsible for IPC production in *D. discoideum*, we set out to generate a *css2* knockout strain. Unfortunately, all our efforts toward this end were unsuccessful. In addition, the extensive restriction-enzyme-mediated insertional mutagenesis (REMI) library of *D. discoideum* (66) does not contain any *css2* mutants. This suggests that, analogous to the IPCS-encoding gene in yeast (67), *Dd*CSS2 is an essential gene.

**Fig. 6.**
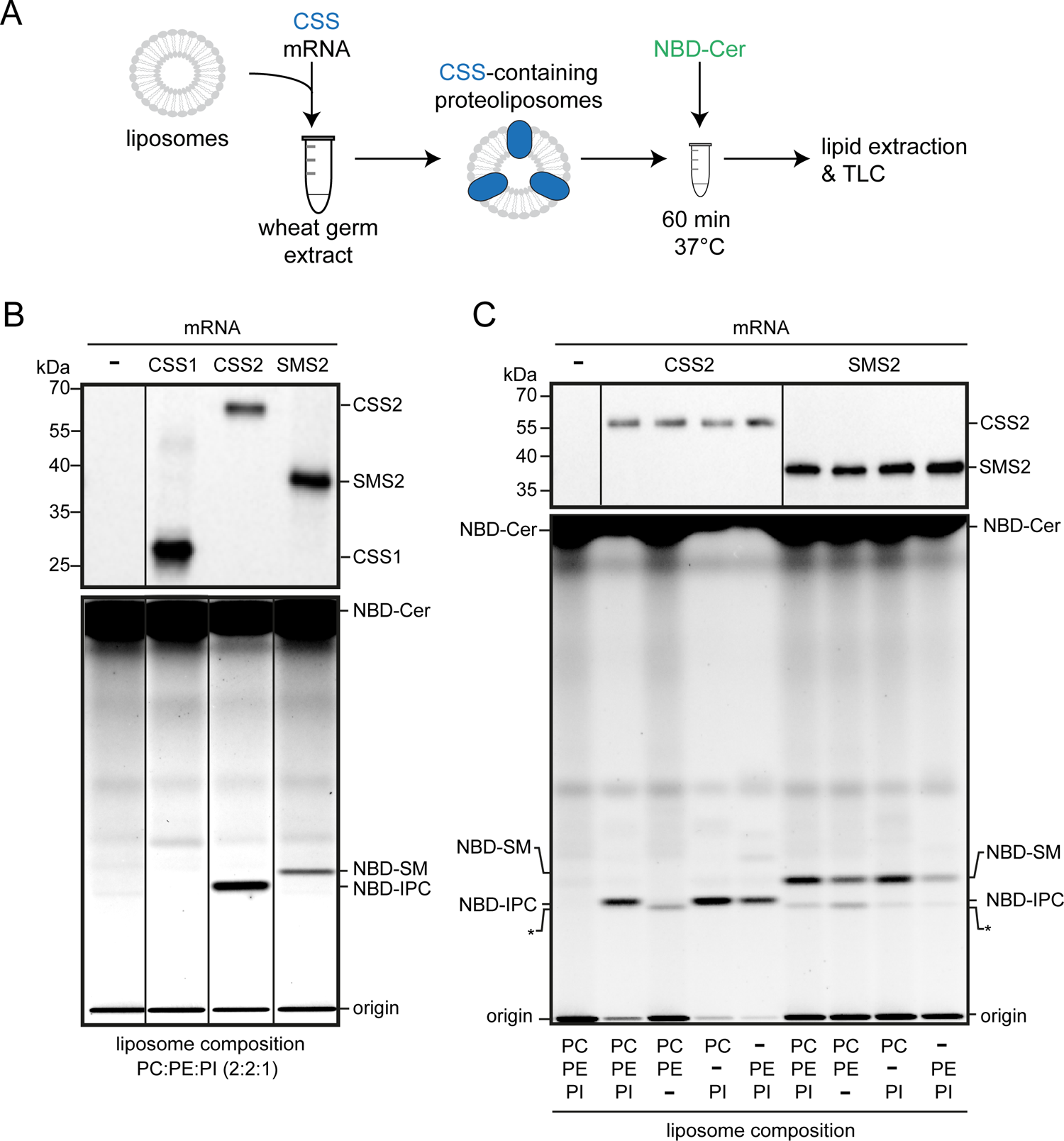
Cell free-expression and functional analysis of *Dd*CSS1 and *Dd*CSS2. A: Schematic outline of the wheat germ-based cell-free translation of CSS mRNA. Unless indicated otherwise, translation reactions were carried out in the presence of liposomes with defined lipid compositions. The resulting proteoliposomes were incubated with NBD-Cer and NBD-labeled reaction products were analysed by TLC. B: Top panel: translation reactions with or without mRNA encoding V5-tagged *Dd*CSS1, *Dd*CSS2 and *Hs*SMS2 carried out in the presence of liposomes with the indicated lipid composition were subjected to western blot analysis using anti-V5 antibody. Bottom panel: TLC analysis of reaction products formed when *Dd*CSS1, *Dd*CSS2 and *Hs*SMS2 produced in the presence of liposomes containing PC:PE:PI (2:2:1) were incubated with NBD-Cer for 60 min at 37°C. C: Top panel: translation reactions with or without mRNA encoding V5-tagged *Dd*CSS2 or *Hs*SMS2 carried out in the presence of liposomes with the indicated lipid composition were subjected to western blot analysis using anti-V5 antibody. Bottom panel: TLC analysis of reaction products formed when *Dd*CSS2 and *Hs*SMS2 produced in the presence of liposomes with the indicated lipid composition were incubated with NBD-Cer for 60 min at 37°C. Migration of an unidentified fluorescent lipid that was present in reactions irrespective of the presence of *Dd*CSS2 or *Hs*SMS2 is marked by an asterisk.

The cyclic depsipeptide antibiotic Aureobasidin A (AbA) is a potent inhibitor of IPC synthase from fungi (68, 69). Indeed, addition of AbA (50 μM) to yeast lysates co-incubated with NBD-Cer and externally added PI effectively blocked the formation of NBD-IPC (Supplemental Fig. S6A). Moreover, the NBD-IPC synthase activity of cell-free produced *Dd*CSS2 was resistant to even high levels of AbA (500 μM). Also, prolonged treatment of *D. discoideum* with AbA had no impact on steady state IPC levels (Supplemental Fig. S6B).

In sum, our results indicate that *Dd*CSS2 functions as an IPC synthase that lacks the AbA sensitivity typical for IPC synthases from fungi. Whether *Dd*CSS2 is required for IPC production in *D. discoideum* remains to be established.

### DdCSS2 localizes to the Golgi apparatus and the contractile vacuole

Previous work revealed that the enzymes responsible for bulk production of IPC, EPC and SM typically localize to the Golgi complex (13, 18, 25). To determine the subcellular localization of *Dd*CSS2, we tagged *Dd*CSS2 with mCherry at either its N-(mCherry-CSS2) or C-terminus (CSS2-mCherry). Expression of the tagged proteins in *D. discoideum* was confirmed by western blotting (Supplemental Fig. S7A). Live cell imaging of these strains by spinning disc (SD) microscopy revealed that both CSS2-mCherry and mCherry-CSS2 displayed a partial co-localization with ZntD-GFP, a zinc transporter present in the juxtanuclear region that is characteristic for the Golgi complex/recycling endosomes in *D. discoideum* (31) (Fig. 7A-B, arrows). The localization of mCherry-tagged *Dd*CSS2 at the Golgi apparatus was confirmed by immunofluorescence (IF) using the Golgi marker AK426 antibody (Fig. 7C-D) (70–72). In contrast, we did not observe any co-localization of mCherry-CSS2 or CSS2-mCherry with the ER marker protein disulfide isomerase (PDI) (Supplemental Fig. S7B). On the other hand, both fusion proteins were present at membrane structures resembling bladders of the CV system prior water expulsion, as well as on vesicle-like structures that were more prevalent in CSS2-mCherry expressing cells (Fig. 8A-B, arrow heads; Supplemental Fig. S7C).

**Fig. 7.**
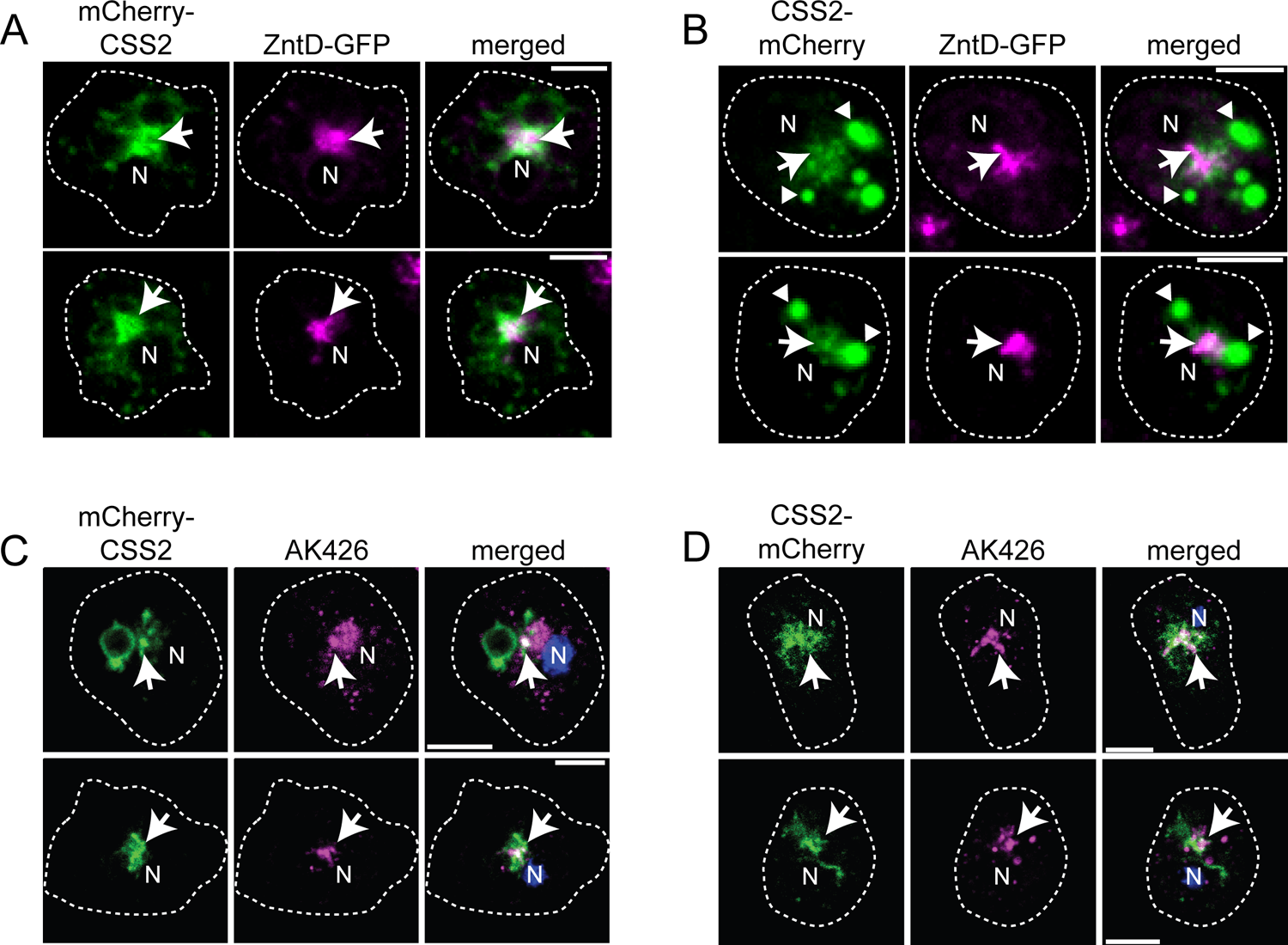
*Dd*CSS2 partially accumulates at the Golgi complex in *D. discoideum*. A and B: CSS2-mCherry as well as mCherry-CSS2 co-localize with ZntD-GFP, a zinc transporter that is located at the Golgi apparatus/recycling endosomes. C and D: CSS2-mCherry as well as mCherry-CSS2 co-localize with the Golgi marker AK426 (71). Cells were either imaged live using SD microscopy (A, B) or fixed with MeOH, stained with the AK426 antibody and imaged using confocal laser scanning microscopy (CLSM) (C, D). Arrows indicate co-localization, arrowheads point to CSS2-mCherry-positive vesicles. N: nucleus. Scale bars, 5 µm.

**Fig. 8.**
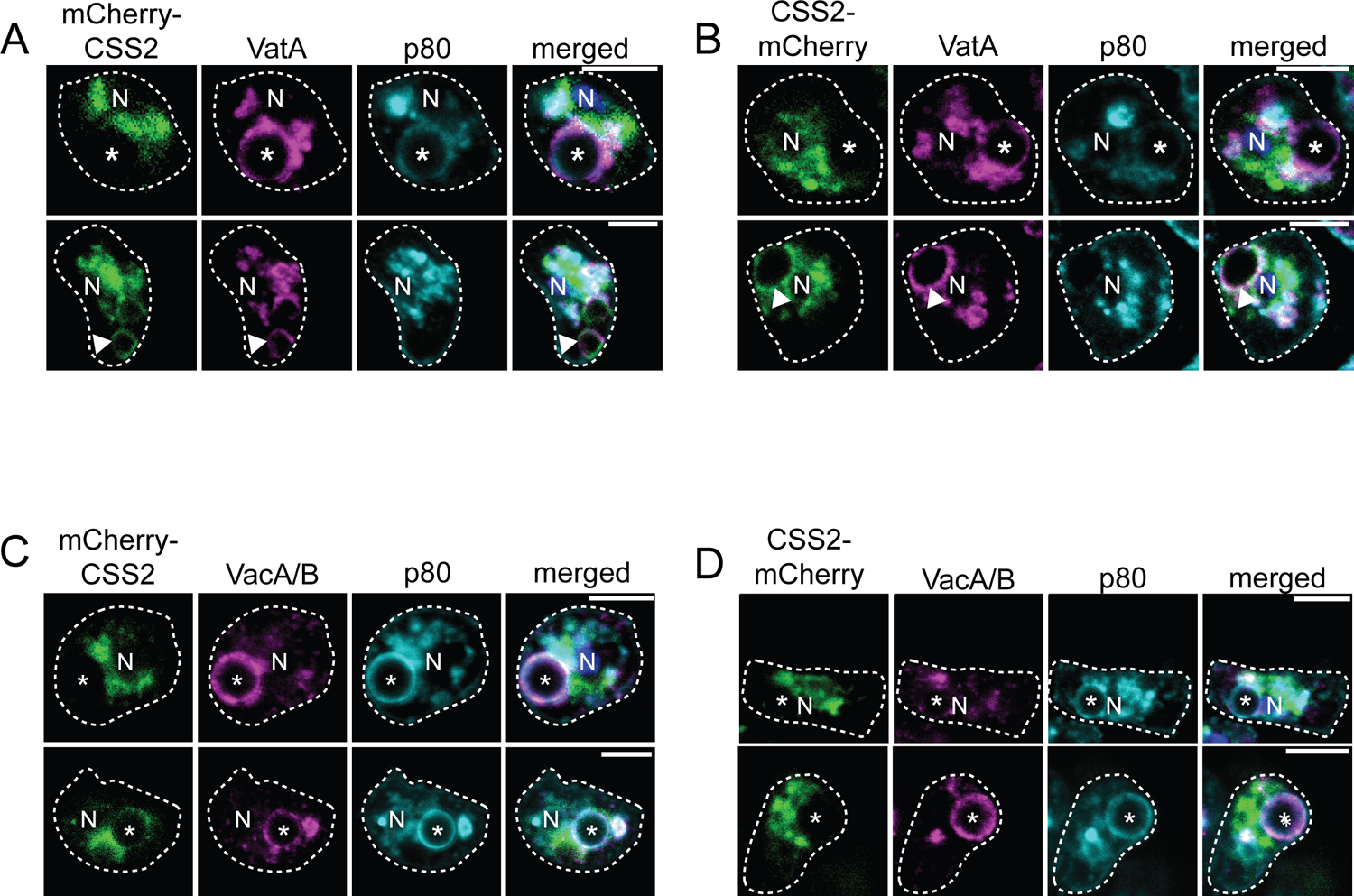
*Dd*CSS2 partially accumulates at the contractile vacuole (CV) in *D. discoideum*. A and B: CSS2-mCherry and mCherry-CSS2 partially co-localizes with H^+^-vATPase-positive, p80-negative structures, indicative of the CV. C and D: CSS2-mCherry and mCherry-CSS2 do not accumulate on BCPs. CSS2-Cherry and mCherry-CSS2 expressing cells were incubated with 3 μm latex beads, fixed with PFA/picric acid (A, B) or MeOH (C, D) and stained with either anti-VatA (A, B) or anti-VacA/B (C, D) and anti-p80 antibodies. Images were generated by CLSM. Asterisks label BCPs. Arrowheads label CSS2-positive CV bladders. N: nucleus. Scale bars, 5 µm.

To analyse the localization of *Dd*CSS2 on CV bladders, cells expressing the mCherry-tagged proteins were fed with latex beads, fixed and then stained with an antibody against VatA, a subunit of the vATPase that is present in both, lysosomes and membranes of the CV network (Fig. 8A-B, arrowheads) (73). To distinguish between the CV and lysosomes, cells were additionally stained for p80, a copper transporter that resides in all endosomes in *D. discoideum* and accumulates on latex beads upon phagosomal maturation (i.e. lysosomes: p80-low; post-lysosomes: p80-high) (74). Indeed, the presence of both mCherry-tagged CSS2 versions at VatA-positive, p80-negative compartments (Fig. 8A-B, arrowheads) and their absence at VatA-positive, bead-containing phagosomes (BCPs) (Fig. 8A-B, asterisks) suggests that *Dd*CSS2 is confined to the CV and does not reside in lysosomes. Moreover, we did not observe any co-localization of CSS2-mCherry or mCherry-CSS2 on Vacuolin-positive BCPs, indicating that *Dd*CSS2 is not a phagosomal protein (Fig. 8C-D).

From this we conclude that the IPC synthase *Dd*CSS2 in *D. discoideum* resides in the Golgi. Moreover, our findings demonstrate an additional localization of the enzyme at the CV.

## Discussion

While *D. discoideum* has been extensively used as experimental model in cell and infection biology, information on its sphingolipidome and the underlying sphingolipid biosynthetic pathway has remained scarce. Combining BLAST searches for homologs of sphingolipid biosynthetic enzymes with LC-MS/MS-based lipid profiling and the functional characterization of cell-free expressed enzymes, we found that *D. discoideum* primarily synthesizes phosphoinositol-containing sphingolipids with phytoceramide backbones. Identification of the corresponding IPC synthase, *Dd*CSS2, revealed a polytopic membrane protein that shares multiple sequence motifs with both yeast IPC synthases and human SM synthases. Interestingly, *Dd*CSS2 displays a dual localization in *D. discoideum*, being present in both the Golgi apparatus and the organism’s osmoregulatory vacuole.

Our untargeted LC-MS/MS lipidome analyses revealed that, in line with previous studies (27, 60), the phospholipid pool in *D. discoideum* is mostly composed of ether lipids, i.e. lipids with ether-linked acyl-chains that are less abundant in mammals (75, 76) and absent in plants or fungi (77). Analogous to the situation in *E. coli* (78) and *D. melanogaster* (79), we found that PE is by far the most abundant phospholipid class in *D. discoideum.* In contrast, PC and PI in each case comprised only a small fraction of the total phospholipid pool. Eukaryotic organisms typically use the most abundant phospholipid as headgroup donor in the production of complex phosphosphingolipids, e.g. PC for the production of SM in mammals (57), PE for the production of EPC in *D. melanogaster* (18, 22), and PI for the production of IPC in plants and yeast (58, 59). Our finding that *D. discoideum* produces IPC in spite of having PE as most abundant phospholipid class provides a significant break in this trend. Analogous to plants and yeast (80), *D. discoideum* mainly synthesizes IPC species containing a phytoceramide backbone with amide-linked acyl chains of 16 to 22 carbon atoms in length. Thus, the fatty-acid moieties in IPC species of *D. discoideum* are significantly shorter than those in yeast. While Cer (40:0;3) is the most abundant Cer in *D. discoideum*, the levels of IPC (40:0;3) are relatively low. This suggests that the IPC synthase in *D. discoideum* may prefer phytoceramides with α-hydroxylated acyl chains as substrate.

IPC species in *D. discoideum* may undergo further head group modifications such as mannosylation for the production of mannosylinositol phosphorylceramide (MIPC) in yeast or glucuronosylation for the production of glycosylinositol phosphorylceramides (GIPC) in plants. While our lipidome analysis so far did not uncover such IPC derivatives, the *D. discoideum* genome encodes a putative homologue of the plant inositol phosphoryl ceramide glucuronosyl transferase 1 (IPUT1) (DDB_G0286945) (Supplemental Table 3), which transfers a glucuronic acid residue to the IPC headgroup in *A. thaliana* (81). Altogether, we conclude that the sphingolipidome of *D. discoideum* is unique and, if any, possesses greater similarities to the sphingolipidomes of *S. cerevisiae* and plants than those of animals.

BLAST searches for homologues of core metabolic enzymes enabled us to partially reconstruct the sphingolipid biosynthetic pathway in *D. discoideum.* However, this strategy did not yield any information on the enzymes responsible for the production of complex sphingolipids. As complementary approach, we searched for polytopic membrane proteins in *D. discoideum* having sequence motifs in common with known IPC and SM synthases, yielding *Ds*CSS1 and *Dd*CSS2. Subsequent analysis of *Ds*CSS1 did not provide any information on its enzymatic activity. However, a BLAST search revealed similarities between *Dd*CSS1 and the yeast diacylglycerol pyrophosphate phosphatase 1 (*Sc*Dpp1), an enzyme that dephosphorylates diacylglycerol pyrophosphate to generate phosphatidate (PA) and subsequently dephosphorylates PA to release diacylglycerol. Whether *Dd*CSS1 mediates dephosphorylation of PA remains to be established.

*Dd*CSS2, on the other hand, qualified as IPC synthase based on the following criteria: (i) cell-free expressed *Dd*CSS2 supports the production of IPC when supplemented with ceramide and PI as headgroup donor; (ii) *Dd*CSS2 shares sequence motifs containing active site residues with known IPC synthases in plants and yeast; (iii) *Dd*CSS2 is predicted to adopt a membrane topology similar to that of IPC synthases in plants and yeast whereby the active site residues are facing the exoplasmic leaflet, hence the side of the membrane where IPC production is thought to occur; (iv) *Dd*CSS2 localizes to the Golgi apparatus, the organelle previously established as the principle site of IPC production in plants and yeast (13, 25, 59, 65). Consequently, we propose to rename *Dd*CSS2 as *Dd*IPCS1 for *D. discoideum* IPC synthase 1.

In spite of its close relationship with the yeast IPC synthase *Sc*Aur1, *Dd*IPCS1 displayed resistance to the cyclic depsipeptide antibiotic AbA, which is a potent inhibitor of the yeast enzyme (20, 23, 82). Inhibition of *Sc*Aur1 by AbA is dependent on residues 137, 157 and 158 (69, 83), which are part of a sequence motif that is not well conserved in *Dd*IPCS1 (Supplemental Fig. S4B, magenta box). AbA was originally isolated from *Aureobasidium pullulans*, a yeast-like fungus found in the natural habitat of *D. discoideum* (84). Thus, it is conceivable that mutations in *Dd*IPC1 enabled *D. discoideum* to develop resistance against the antibiotic activity of AbA. Also of note is the dual location of *Dd*IPC1, which resides in both the Golgi apparatus and the CV. In *D. discoideum* and other free-living organisms, the CV is an osmoregulatory organelle for water expulsion under hypotonic conditions (85). Water discharge is carried out by a giant kiss-and-run exocytic event. CV discharge may provide an alternative mode to deliver IPC produced by *Dd*IPCS1 to the PM. This may be necessary to sustain the gradient of complex sphingolipids along the secretary pathway, with the highest concentration at the PM.

Whether *Dd*IPCS1 is the sole IPC synthase in *D. discoideum* remains to be established. All our efforts to generate *Dd*IPCS1knockouts by homologues recombination were unsuccessful, suggesting that *Dd*IPCS1 is essential for *D. discoideum* growth and survival. Similar observations have been made for IPC synthases in *A. thaliana* and *S. cerevisiae* (69, 81) while in *T. brucei*, RNAi-mediated knock-down of the entire gene locus encoding *Tb*SLS1-4 is lethal (86). Future *Dd*IPCS1 knockdown studies should help elucidate the biological roles of IPC production in *D. discoideum* in particular during host-pathogen interactions. Given that *D. discoideum* was exploited by pathogens well before the evolution of mammals, we anticipate that such studies may uncover very fundamental sphingolipid-dependent mechanisms underlying infection.

## Data availability statement

All data are included in the manuscript and supporting information. In addition, we provide a lipidomics minimal reporting checklist from the Lipidomics Standard Initiative (Supplemental Table 4).

## Acknowledgments

We greatly acknowledge the integrated Bioimaging facility (iBiOs) at the University of Osnabrück and especially Rainer Kurre for his expertise and friendly support. We thank Thierry Soldati, Dominik Schwudke and Nicolas Gisch for carefully reading this manuscript and thoughtful suggestions. This work was supported by the Deutsche Forschungsgemeinschaft (SFB1557-P1 to C.B. and SFB1557-P7 to J.C.M.H). The Barisch lab is a member of the SPP2225.

## Conflict of interests

The authors declare that they have no conflict of interest.

## Supporting information

**Supplemental Fig. S1.**
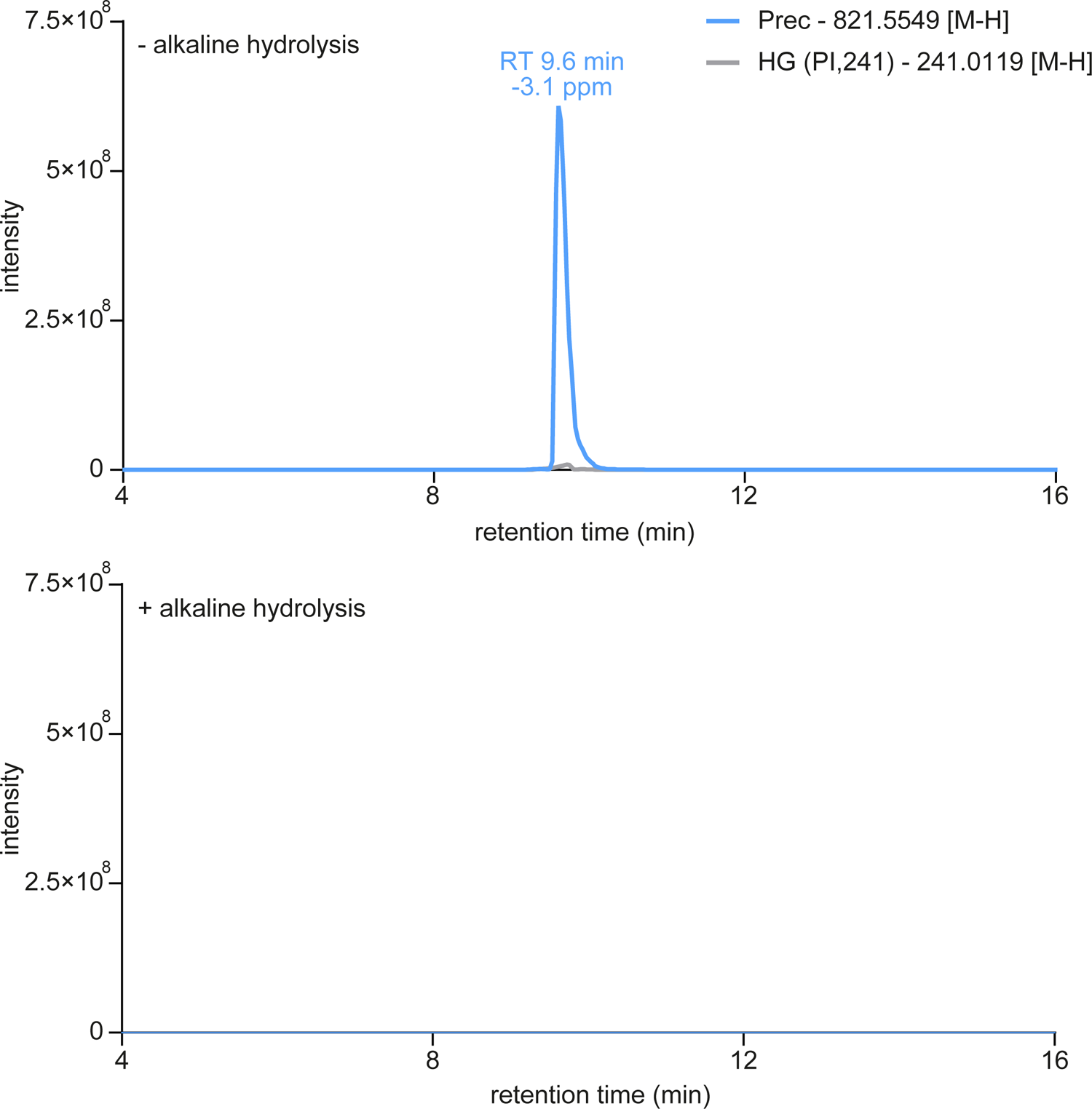
Deacylation of the phospholipids by alkaline hydrolysis. Intensity plots of PI (O-34:1) of samples subjected (+) or not (-) subjected to alkaline hydrolysis. *D. discoideum* was grown in HL5c (complex) medium. Lipids were extracted following (34) (+ alkaline hydrolysis) and (32) (-alkaline hydrolysis).

**Supplemental Fig. S2.**
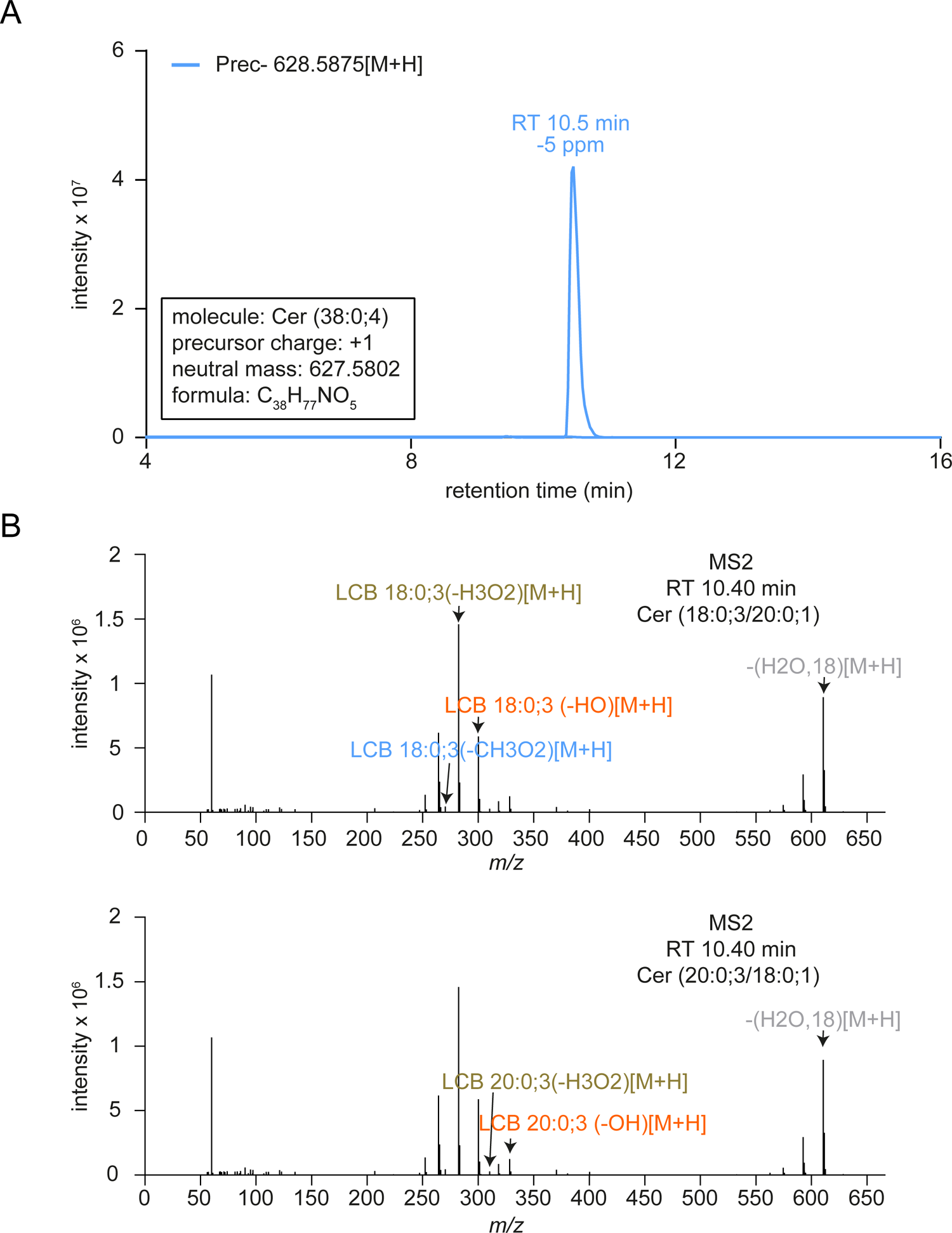
Resolving the LCB chain length of ceramide (Cer). A: Precursor chromatogram of Cer (38:0;4). B: Product ion fragmentation pattern of Cer (38:0;4) including annotations of Cer (18:0;3/20:0;1) (top) and Cer (20:0;3/18:0;1) (bottom) fragments. *D. discoideum* was grown in HL5c (complex) medium. Lipids were extracted following (34) with alkaline hydrolysis.

**Supplemental Fig. S3.**
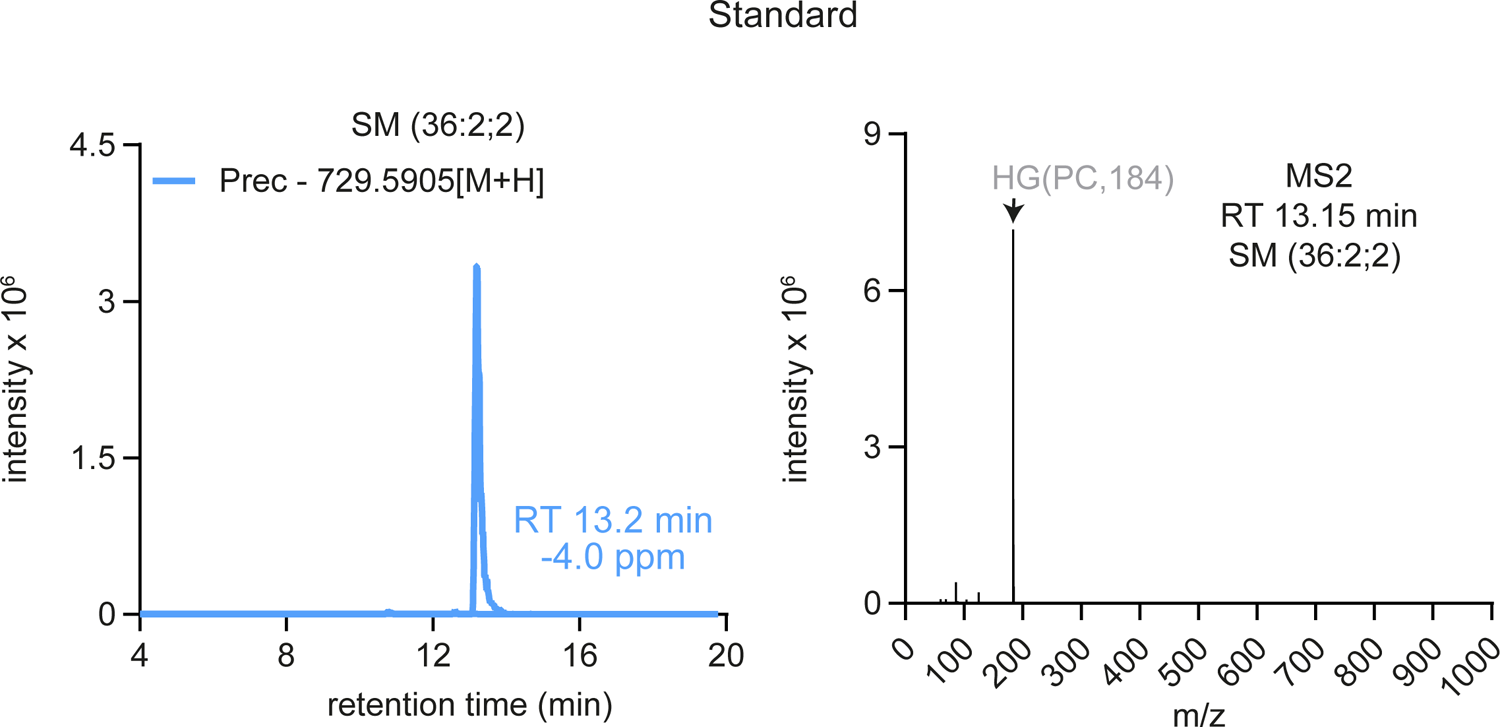
Intensity plot of SM. SM (36:2;2) from a commercial standard (Avanti Polar Lipids) served as positive control for SM detection.

**Supplemental Fig. S4.**
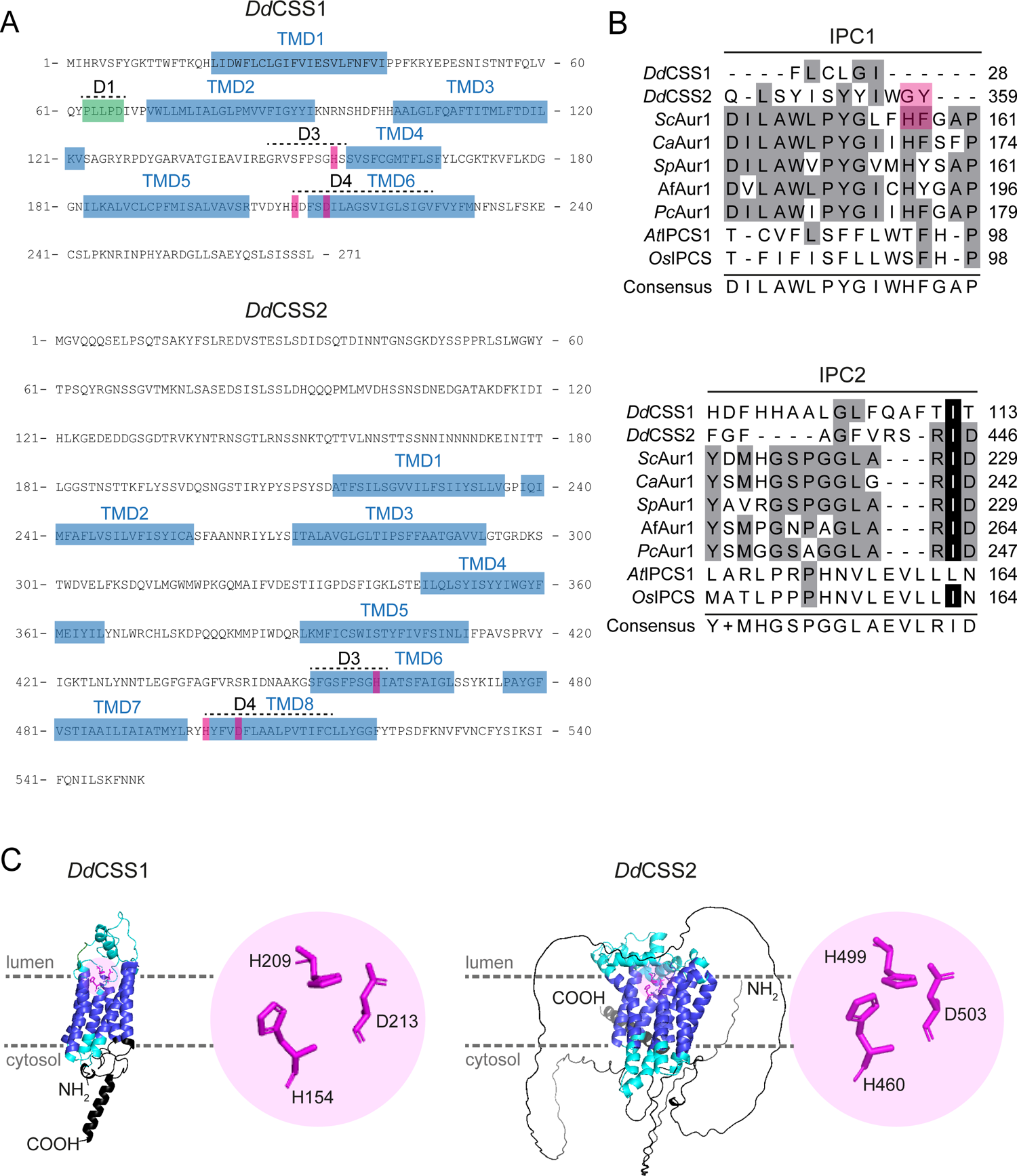
Conserved sequence motifs in *Dd*CSS1 and *Dd*CSS2. A: Amino acid sequences of *Dd*CSS1 and *Dd*CSS2. Regions predicted to form transmembrane domains (TMD1-TMD8) are highlighted in blue. Amino acids of the catalytic triad are marked in magenta, a potential SMS D1 motif is labelled in green. The sequences that are highly similar to D1, D3 and D4 motifs from human SM synthases are indicated. B: Alignment of two domains conserved in fungal IPC synthases (named IPC1 and IPC2). Black indicates identical amino acids, grey conservative amino acid substitutions. Residues that confer AbA resistance in *Sc*Aur1 and the corresponding amino acids in *Dd*CSS2 are highlighted in magenta. Alignment was performed with Jalview workbench (87), using the in-built MAFFT alignment option (G-INS-i). C: AlphaFold-derived structures of *Dd*CSS1 and *Dd*CSS2. The inset shows the amino acids composing the catalytic triad (magenta). Luminal- and cytosolic-exposed regions are indicated in cyan, TMDs are presented in violet.

**Supplemental Fig. S5.**
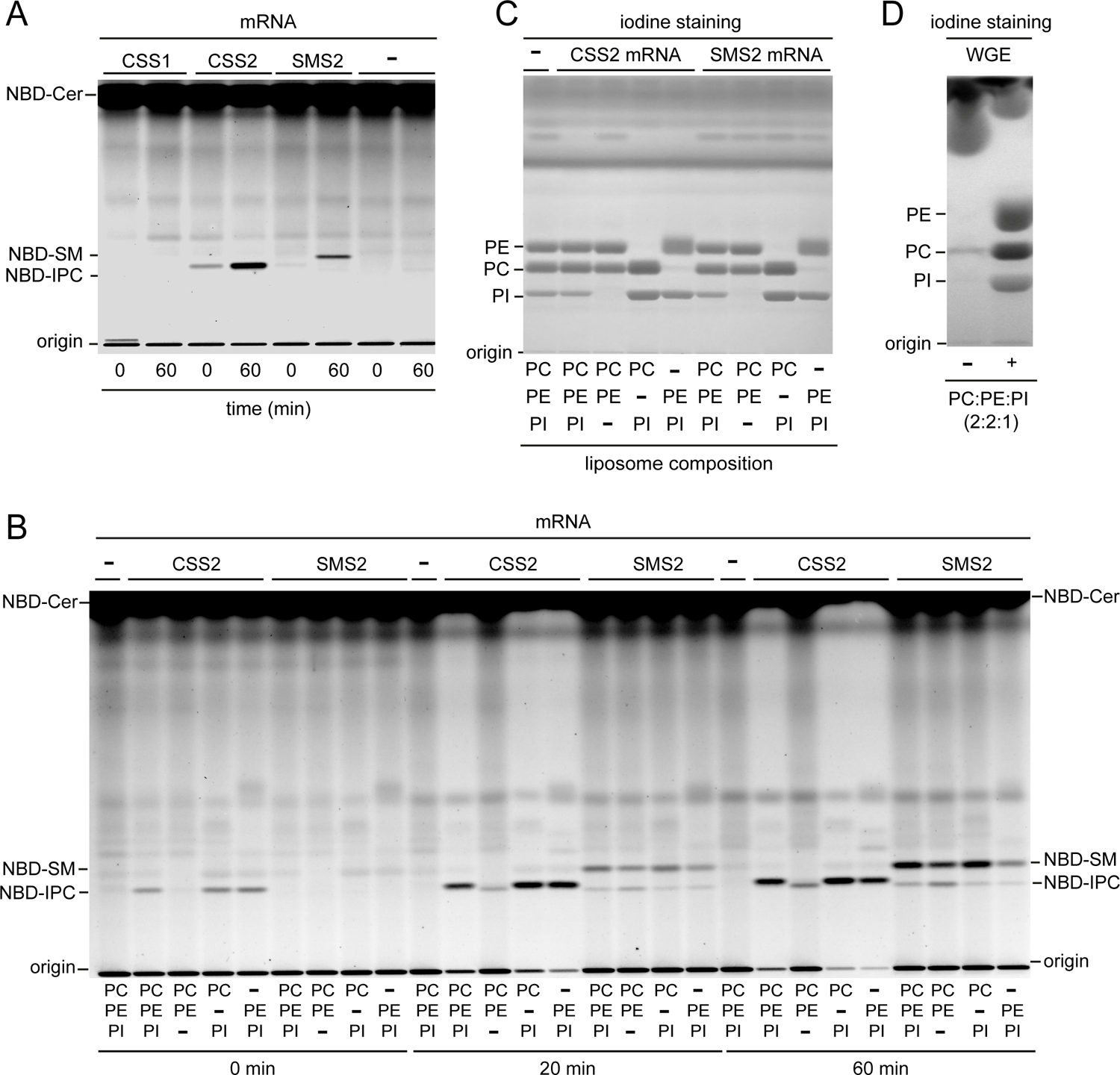
Complete TLCs and iodine staining. A: TLC analysis of reaction products formed when the translational reactions of *Hs*SMS2, *Dd*CSS1 and *Dd*CSS2 were incubated with liposomes containing PC:PE:PI (2:2:1). B: TLC analysis of reaction products of *Hs*SMS2 and *Dd*CSS2 that were incubated with liposomes of varying phospholipid composition. The reaction was stopped at the indicated time points. C: Iodine staining of B. Shown are the reaction products after 60 min incubation. D: Iodine staining of the WGE with and without the addition of liposomes (PC:PE:PI; 2:2:1).

**Supplemental Fig. S6.**
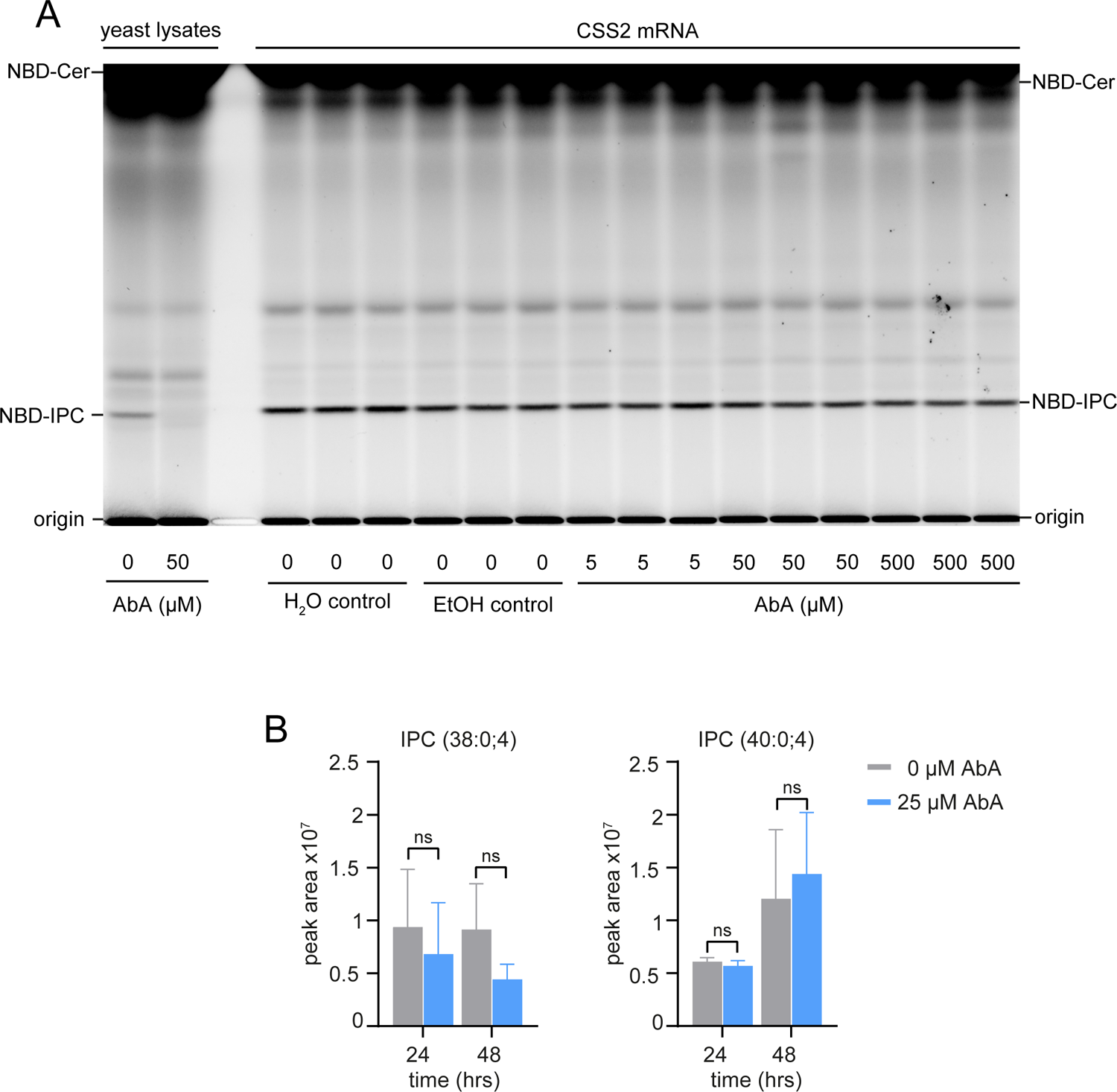
Analysis of IPC synthesis by Aureobasidin (AbA). A: TLC analysis of reaction products of *Dd*CSS2 that were incubated with AbA. Yeast lysates incubated with AbA served as a positive control. B: LC/MS-MS analysis of IPC(38:0,4) and IPC(40:0,4) after treatment with AbA. HL5c grown-*D. discoideum* was treated with or without 25 μM of AbA for 24 or 48 hrs, prior to extraction by (32). Graph shows mean and ±SD (n=4). Statistics were assessed with a paired t-test. ns: not significant.

**Supplemental Fig. S7.**
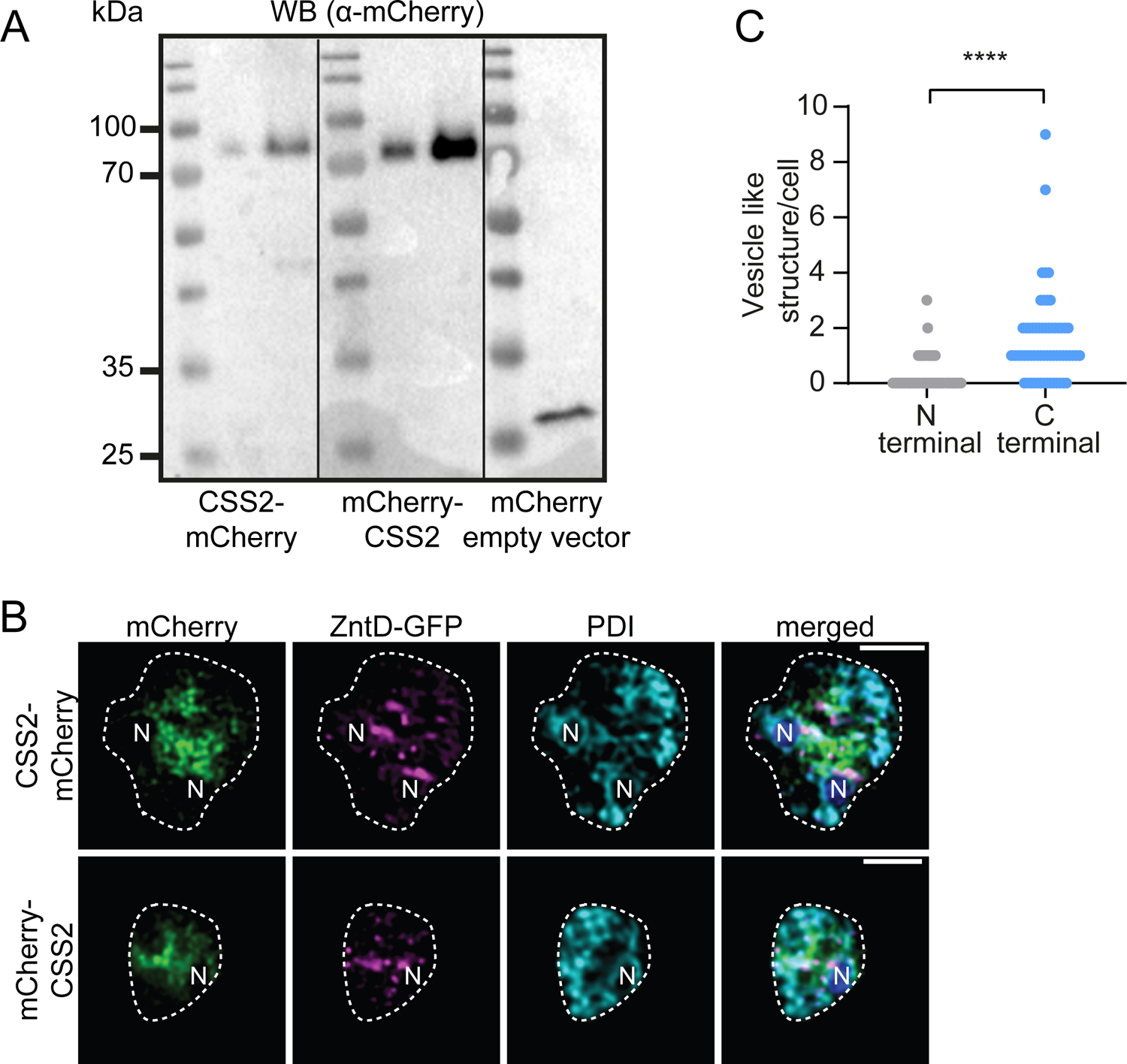
Expression of fusion constructs and localization analysis. A: Western blot of mCherry-CSS2 and CSS2-mCherry with untagged mCherry as a control. 2×10^4^ and 10×10^5^ cells were used for the first and second lanes, respectively. B: CSS2-mCherry and mCherry-CSS2 do not co-localize with the ER-marker PDI. Cells were fixed with MeOH, stained for PDI and imaged by CLSM. Images were deconvolved. N: nucleus. Scale bars, 5 µm. C: Quantification of CSS2-mCherry and mCherry-positive vesicles. Statistical differences were calculated with an unpaired t-test (**** p < 0.0001). Error bars indicate ±SD (n=3; number of cells = 60).

**Supplemental Table 1.** Material used within this publication.

**Supplemental Table 2.** Complete dataset of the lipidomics and sphingolipidomics analysis described in Fig. 2 and Fig. 3.

**Supplemental Table 3.** Excel file including the results of the comprehensive BLAST search to identify *D. discoideum* homologues of sphingolipid synthesising enzymes. BLAST searches were carried out using the BLASTp function from Dictybase (http://dictybase.org/tools/blast) with default option. Sequences from *H. sapiens*, *S. cerevisiae* and *A. thaliana* were derived from Uniprot and used as baits.

**Supplemental Table 4.** Lipidomics minimal reporting checklist from the Lipidomics Standard Initiative (https://lipidomicstandards.org/).

## Notes

### Competing Interest Statement

The authors have declared no competing interest.

### Summary of Updates

Typos have been corrected.

